# Deepurify: a multi-modal deep language model to remove contamination from metagenome-assembled genomes

**DOI:** 10.1101/2023.09.27.559668

**Authors:** Bohao Zou, Jingjing Wang, Yi Ding, Zhenmiao Zhang, Yufen Huang, Xiaodong Fang, Ka Chun Cheung, Simon See, Lu Zhang

## Abstract

Metagenome-assembled genomes (MAGs) offer valuable insights into the exploration of microbial dark matter using metagenomic sequencing data. However, there is a growing concern that contamination in MAGs may significantly impact the downstream analysis results. Existing MAG decontamination methods heavily rely on marker genes but do not fully leverage genomic sequences. To address the limitations, we have introduced a novel decontamination approach named Deepurify, which utilizes a multi-modal deep language model employing contrastive learning to learn taxonomic similarities of genomic sequences. Deepurify utilizes inferred taxonomic lineages to guide the allocation of contigs into a MAG-separated tree and employs a tree traversal strategy for maximizing the total number of medium- and high-quality MAGs. Extensive experiments were conducted on two simulated datasets, CAMI I, and human gut metagenomic sequencing data. These results demonstrate that Deepurify significantly outperforms other decontamination methods.

## Introduction

Short-read metagenomic sequencing has gained popularity in investigating unculturable microbial genomes [1, 2, 3, 4], but single contigs assembled by short-reads often lead to fragmented and incomplete microbial genomes [5, 6, 7]. Several contig binning tools [8, 9, 10, 11] have been developed to group contigs into metagenome-assembled genomes (MAGs) based on their abundances and sequence contexts to represent microbial genomes. Several studies [12, 13, 14] claimed the qualities of those MAGs were comparable to the genomes from microbial isolates, but there has been a growing concern that contamination may seriously impact the qualities of MAGs [15]. MAG contamination refers to a mixture of contigs from different microbes in the same MAG and those chimeric MAGs would substantially reduce the reliability of downstream ecological and evolutionary analyses. Bowers et al. [16] suggested eliminating the MAGs with more than 10% contamination, but many microbes from MAGs with marginal contamination would be missed. In our preliminary study, we observed a considerable number of MAGs would be removed due to their marginal contamination values, even for some high-abundance MAGs (**Supplementary Note** 1). This may result in the loss of a significant number of MAGs for subsequent downstream analysis.

Several tools [17, 18, 19, 20] have been developed to identify and remove the potentially contaminated contigs from chimeric MAGs based on marker genes and the sequence characteristics from known species. Two pipelines [17, 18] published several years ago are no longer actively supported and have not been widely accepted by the community. More recent and actively supported tools are MAGpurify [19] and MDMcleaner [20]. MAGpurify was recently developed for MAG decontamination using three sources of information: the phylogenetic or clade-specific marker genes, the GC contents, and tetranucleotide frequencies of contigs. MDMcleaner utilizes marker genes (coding, 16S, and 23S rRNA genes) to predict the taxonomic classification of contigs. The contig taxonomies are determined by the taxonomic Least Common Ancestor (LCA) of the involved marker genes and any contigs that have different annotations with the dominating taxon of the MAG would be removed.

Although MAGpurify and MDMcleaner show promising results, they suffer from several issues that hinder their widespread applications in various scenarios. First, both of them need to align marker genes/contigs to the reference databases and this approach is inapplicable to novel microorganisms. As previously observed, it has been noted that the reference genomes available in RefSeq (117,030 as of March 11, 2022) only account for less than 5.319% of all species [21]. In addition, the alignment is time-consuming even if the built-in databases are optimized (**Supplementary Note** 2). Second, previous study [22] has pointed out that many factors have the potential to reduce the performance of alignment-based tools on phylogenetic analysis, such as sequence misalignment, false-orthologous assignment, gene duplication or loss events, horizontal gene transfer, and the presence of homoplasy, etc. Third, various genomic alterations, including genomic variations, alterations in gene order, and genome rearrangements, among others, have been identified as factors enhancing the resolution and reliability of differentiating the genomic sequences from different species [23]. These forms of evidence can offer invaluable insights that are uniquely attainable through whole genome sequences. Fourth, we found a majority of contamination in MAGs occurred at the genus and species levels (**Supplementary Note** 3) and both MAGpurify and MDMcleaner demonstrated poor performance at these low taxonomic ranks (**Supplementary Note** 4).

In this study, we developed Deepurify for MAG decontamination with high resolution and generalization using a multi-modality deep language model. In the training procedure, Deepurify developed two distinct encoders, a genomic sequence encoder (GseqFormer, **Methods**) and a taxonomic encoder (Long short-term memory, LSTM) to encode genomic sequences and their source genomes’ taxonomic lineages, respectively. Next, Deepurify learned their relationships in different taxonomic ranks using contrastive training (Figure 1). In the decontamination process, Deepurify initially quantified the taxonomic similarities of contigs by assigning taxonomic lineages to them (Figure 2 **a**). It then used these lineages to construct a MAG-separated tree, partitioning the MAG into distinct sections, each containing contigs with the same lineage (Figure 2 **c**). This approach optimized contig utilization within the MAG, avoiding immediate removal of contaminated contigs. It was especially effective for MAGs with high contamination rates. Lastly, a tree traversal algorithm was devised to maximize the count of medium- and high-quality MAGs within the MAG-separated tree (Figure 2 **d**).

**Figure 1:**
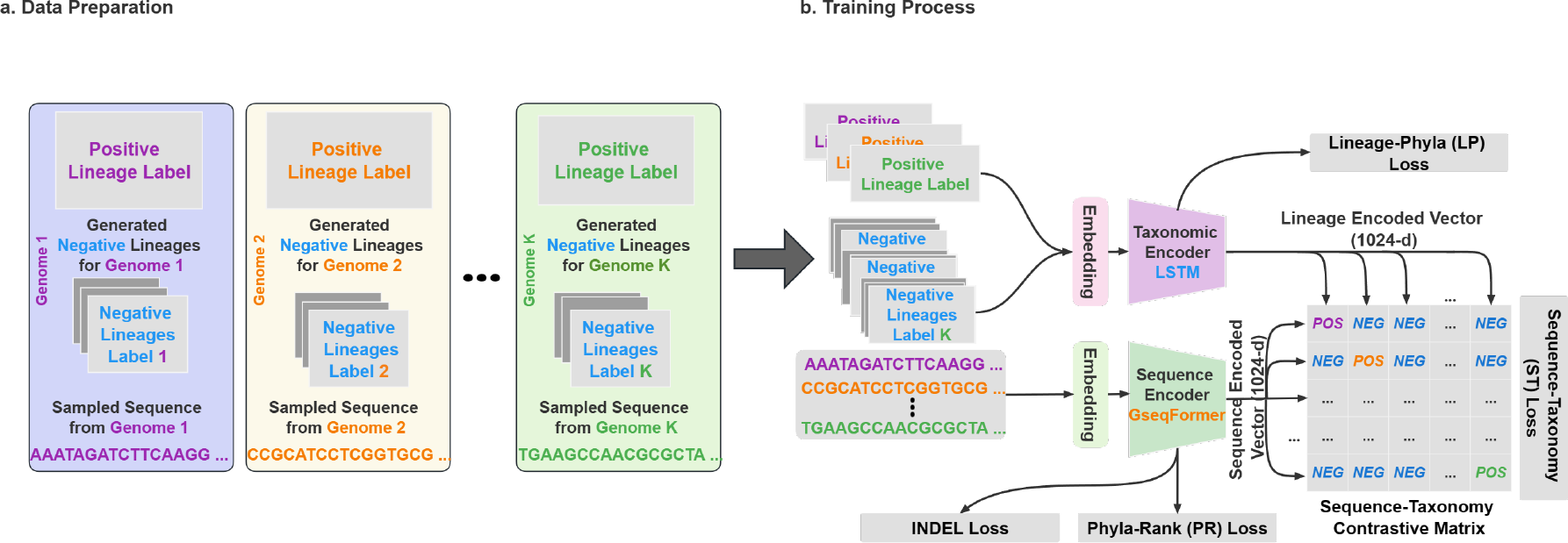
The Deepurify training procedure consisted of two phases: data preparation and the training process. (**a**). In data preparation, Deepurify used the positive lineage label of each genome, generated multiple negative lineage labels for each genome, and sampled an appropriate-length sequence from the corresponding genome. (**b**). During training, the taxonomic encoder encoded positive and negative lineage labels. Sequences were encoded using GseqFormer. A sequence-taxonomy contrastive matrix was built based on calculating the cosine similarity between encoded sequences and lineages. The cosine similarity between the positive label and the sequence is anticipated to surpass that between negative labels and the sequence. Therefore, the ST loss accounted for the majority of the training losses, whereas the other losses facilitated the training process and improved the model’s robustness.

**Figure 2:**
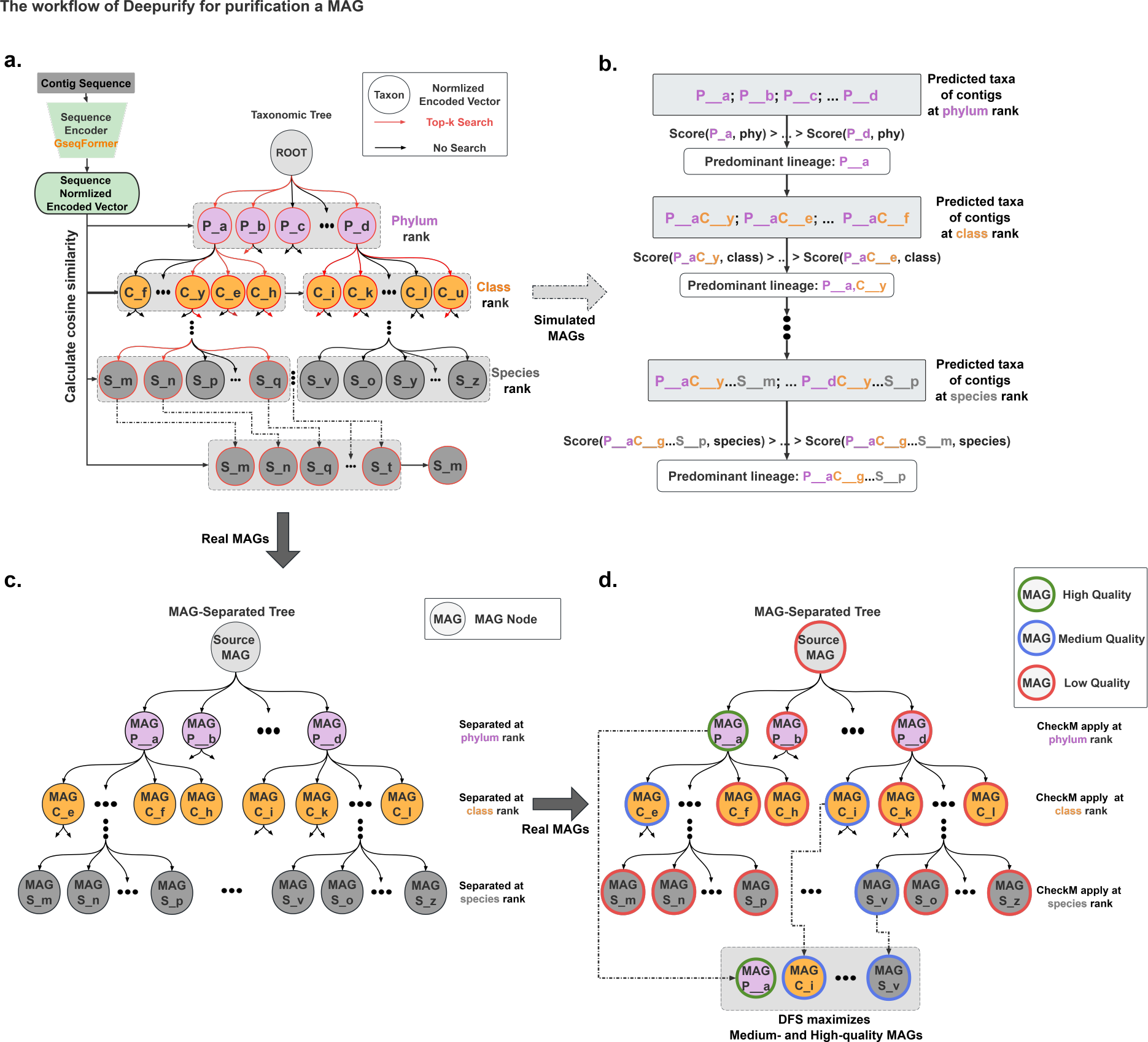
The purification workflow of Deepurify. (**a**). Deepurify assesses taxonomic similarities among sequences through the assignment of taxonomic lineages. It employs a top-k search approach within the taxonomic tree to identify candidate lineages, subsequently selecting the lineage with the highest similarity to the sequences. (**b**). Deepurify applies a scoring function to the lineage of contigs to determine the predominant lineage of contigs in the MAG. The taxon with the highest score is chosen as the predominant lineage at different ranks. This process crosses ranks from phylum to species, ensuring the predominant lineage is consistent and coherent. (**c**). For optimal contig utilization within a MAG without dropping contaminated contigs directly, Deepurify constructs a MAG-separated tree. This tree partitions the MAG based on predicted lineage. Each node contains contigs sharing the same taxon at that rank. To prevent duplicate single-copy genes (SCGs), Deepurify applies SCGs to each node. (**d**). Deepurify employs a depth-first search (DFS) algorithm on the MAG-separated tree to maximize the total number of high- and medium-quality MAGs.

We observed that Deepurify outperformed two state-of-the-art tools MAGpurify and MDMcleaner in simulated, CAMI I challenge (high, medium1, medium2, and low) [24] and human gut metagenomic sequencing data [25, 26]. For simulated data, chimeric MAGs were created by mixing the sequences from two microbial genomes at various taxonomic ranks, with contamination rates from 5% to 20%. Deepurify achieved balanced macro F1-scores almost twice as high as that of MAGpurify across all taxonomic ranks and 1.5 times higher than that of MDMcleaner at the genus and species ranks (Figure 3, **Supplementary Table** 3) on average. Additionally, Deepurify demonstrated outstanding generalization capabilities, where it achieved excellent accuracy in identifying contaminated contigs even if their source genomes were absent from the training set (Figure 4, **Supplementary Table** 4). For CAMI I and a human gut metagenomic sequencing dataset, *S*1 [25], we applied Deepurify to the results of four mainstream contig binning tools (VAMB [8], CONCOCT [9], MetaBAT2 [11], and MaxBin [10]), and the results showed that it could substantially improve MAG quality, surpassing both the MAGpurify and MDMcleaner for all binning tools. Next, we applied Deepurify to a large metagenomic sequencing dataset derived from a diarrhea-predominant Irritable Bowel Syndrome (IBS-D) cohort, including 290 patients and 89 healthy controls [26]. We found Deepurify could rescue 70.12% highly contaminated MAGs (completeness ≥ 50% and contamination ≥ 25%) to medium-(completeness ≥ 50% and contamination ≤ 10%) and high-quality (completeness ≥ 90% and contamination ≤ 5%) MAGs. The corresponding percentages of MAGpurify and MDMcleaner were only 1.4% and 0.7%, respectively. Moreover, we compared the annotation of these MAGs before and after MAG decontamination and identified five new species (**Supplementary Table** 5) and one new genus (**Supplementary Table** 6). Among them, one of the species demonstrated a suggestive association with IBS-D.

**Figure 3:**
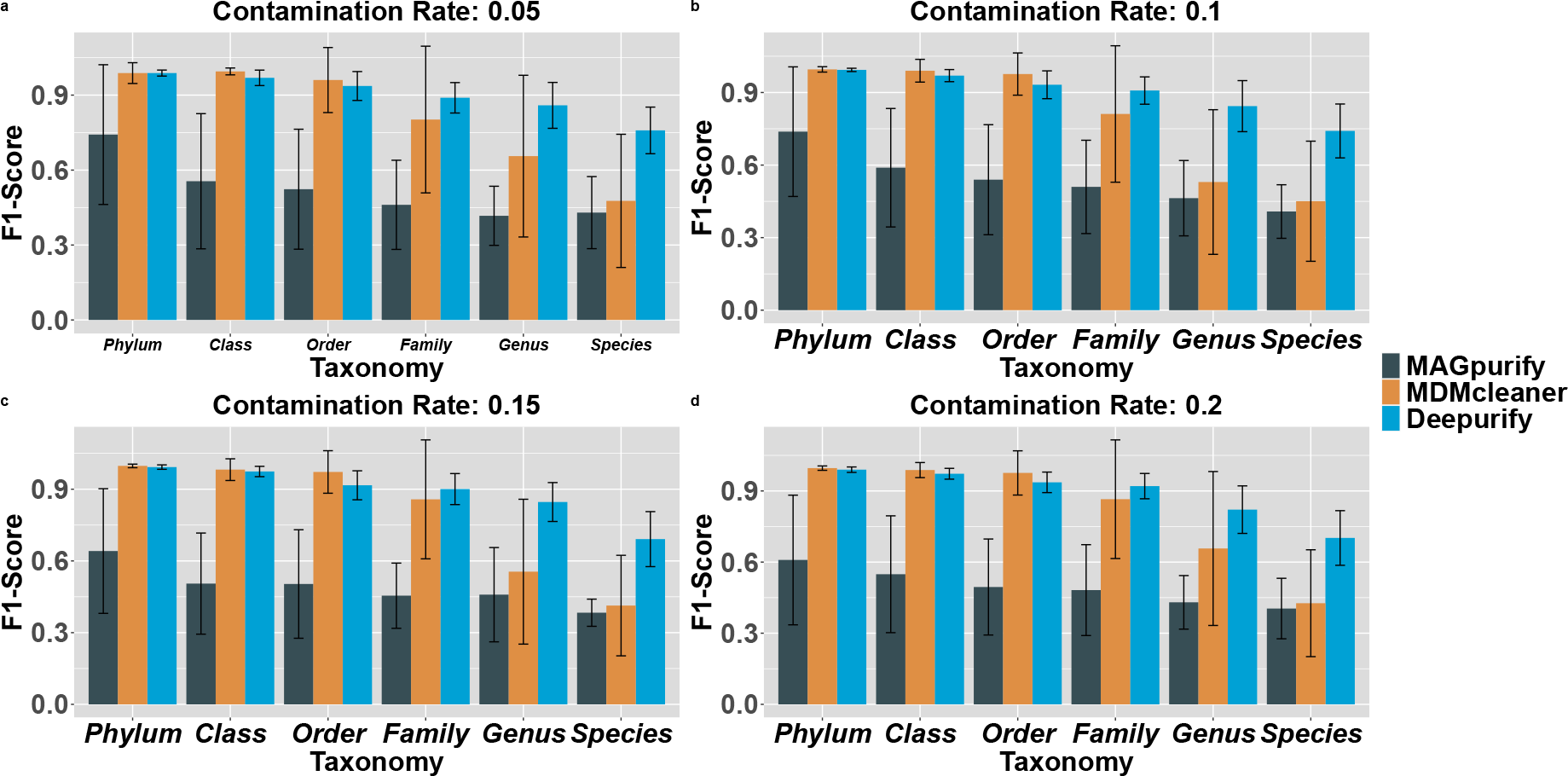
The averaged macro F1-score at various contamination ratios and taxonomic ranks for MAGpurify, MDMcleaner, and Deepurify. The error bars represent standard deviations.

**Figure 4:**
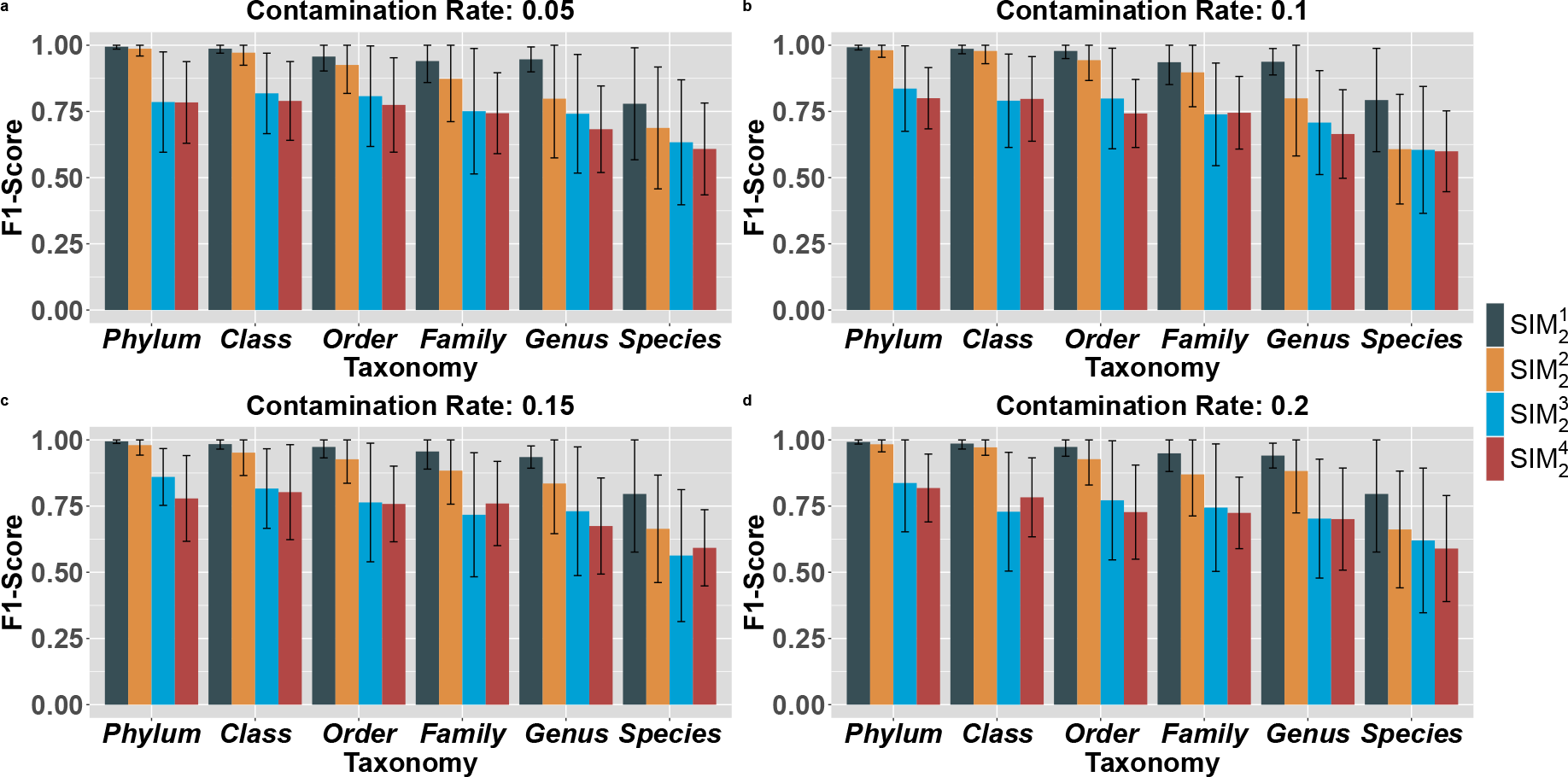
The averaged macro F1-score calculated for 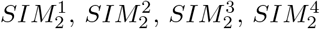 at different contamination ratios and taxonomic ranks for Deepurify.

## Results

### Deepurify architecture and decontamination workflow

Deepurify was a multi-modal deep language model developed specifically to remove contaminated contigs from a MAG. Figure 1 **b** and Figure 2 depict the fundamental architecture and decontamination workflow of Deepurify. Its architecture resembles that of CLIP [27], a well-established multi-modal model incorporating two encoders designed to process data from two modalities: 1). GseqFormer, for encoding genomic sequences, and 2). LSTM, for encoding taxonomic lineages (**Methods**). During training, we utilized contrastive learning to empower Deepurify to distinguish between real (positive) and fake (negative) taxonomic lineages of a sequence (Figure 1). This distinction is based on the cosine similarity between normalized encoded sequences and both positive and negative normalized encoded lineage vectors. Positive encoded lineages should exhibit higher cosine similarity with encoded sequences compared to negative ones. During the decontamination process (Figure 2), Deepurify first assessed the taxonomic similarities of contigs by computing cosine similarity scores between the contigs and lineages in the taxonomic tree. Subsequently, it assigned the lineages to the contigs based on the highest similarity (Figure 2 **a**). Deepurify devised a scheme involving the construction of a MAG-separated tree to maximize the effective utilization of contigs, without directly discarding contaminated ones within a MAG (Figure 2 **c**). This scheme was especially valuable in MAGs with high contamination rates. The MAG-separated tree partitioned contigs within the MAG into distinct branches according to their predicted taxonomic lineages across multiple taxonomic ranks. Each node in the tree contains contigs sharing the same taxon at that rank. Deepurify identified and applied single-copy genes (SCGs) to each node to prevent duplication of SCGs within it. Finally, Deepurify applied CheckM [28] to each node of the tree and employed a depth-first search (DFS) algorithm to traverse the MAG-separated tree to maximize the count of high- and medium-quality MAGs (**Methods**; Figure 2 **d**).

### Development of simulated testing sets

We generated two simulated testing sets, *SIM*_1_ and *SIM*_2_, to evaluate Deepurify’s capability in distinguishing between core and contaminated contigs within a chimeric MAG. The *SIM*_1_ testing set assessed Deepurify’s decontamination performance when the source genomes of both core and contaminated contigs were part of the training set. Conversely, the *SIM*_2_ testing set evaluated its performance when the source genomes of the contigs were either included or excluded from the training set. A simulated chimeric MAG primarily consisted of core contigs, with a minority being contaminated. We referred to the source genomes of core contigs as “core” genomes and the source genomes of contaminated contigs as “contaminated” genomes.

In our chimeric MAG simulation, we simulated contamination occurring at different taxonomic ranks by randomly selecting core and contaminated genomes from two species at varying taxonomic distances on the taxonomic tree. The LCAs of these two species’ lineages ranged from kingdom to genus (lineages differ starting from phylum to species). Each simulated MAG consisted of 200 contigs, with lengths distributed uniformly between 1,000 bps and 8,192 bps. We generated 50 simulated MAGs for different contamination proportions (5%, 10%, 15%, and 20%) at each taxonomic rank of LCA.

The test set of *SIM*_1_ was generated using the genomes that were all included in its training set *GS*_*c*_ (**Methods**). For *SIM*_2_, its training set *GS*_*p*_ (**Methods**) lacked either core or contaminated genomes, resulting in four scenarios for simulation: 1. both core and contaminated genomes included in the *GS*_*p*_ (*SIM* ^1^); 2. only core genomes included in the 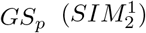 3. only contaminated genomes included in the 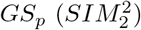 4. both core and contaminated genomes were not included the 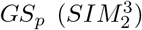. To address the imbalance issue between the number of core and contaminated contigs in a simulated MAG, we utilized a balanced macro F1-score to evaluate the performance of MAGpurify, MDMcleaner, and Deepurify.

### Deepurify has superior purification performance on *SIM*_1_

We applied MAGpurify, MDMcleaner, and Deepurify to *SIM*_1_ testing set and we observed that Deepurify outperformed MAGpurify significantly across all taxonomic ranks and contamination proportions (Figure 3, **Supplementary Table** 3). Compared to MAGpurify, Deepurify increased the overall averaged F1-score by 45.18% (phylum), 76.75% (class), 80.53% (order), 89.75% (family), 90.51% (genus), and 78.02% (species) across different contamination proportions. We observed that Deepurify and MDMcleaner performed comparably when the lineages of core and contaminated genomes differed at higher taxonomic ranks such as phylum, class, and order. However, Deepurify exhibited significant improvements when the differences in lineages began at the family, genus, and species ranks, with an overall average F1-score increase of 8.45% (family), 40.54% (genus), and 63.72% (species) compared to MDMcleaner. This fact suggested Deepurify could be more efficient to be applied in real metagenomic sequencing data, as most of the MAG contamination was found to exist at the genus and species (**Supplementary Note** 3). We noticed that the F1-scores of MAGpurify, Deepurify, and MDMcleaner decreased as the taxonomic ranks became lower. This could be due to the higher proportion of homologous sequences between the core and contaminated genomes at the genus and species taxonomic ranks.

We also observed the standard deviations (SD) of the F1-scores of Deepurify were considerably lower than those of MAGpurify and MDMcleaner suggesting Deepurify was more robust regardless of the sources of contamination. On the one hand, the SD of F1-scores of MAGpurify were consistently reduced at taxonomic ranks from high to low, revealing it is more conservative to remove contigs at low taxonomic ranks. Consequently, it may not effectively remove contaminated contigs when contamination occurs at these lower taxonomic ranks. On the other hand, the SD of F1-scores of MDMcleaner were the highest at genus and species ranks, indicating that it was not stable in accurately distinguishing between genomes with homologous sequences. Furthermore, we observed an opposite trend between the contamination rates and the average F1-score of MAGpurify. This indicates that MAGpurify was not able to eliminate contaminated contigs at high rates of contamination efficiently. Although MDMcleaner’s performance remained relatively stable across different contamination rates, it experienced a significant decline as taxonomic ranks decreased. Deepurify emerged as the most efficient and robust model across all tested conditions.

### Deepurify has strong generalization ability for novel microbes

As MAGpurify and MDMcleaner did not provide any interface to allow users to rebuild their databases, we could not evaluate them on the *SIM*_2_ testing set. We applied Deepurify on *SIM*_2_ and we found that the F1-scores of Deepurify were only marginally reduced regardless of core or contamination genomes absent from the training sets (Figure 4, **Supplementary Table** 4). We used the performance of Deepurify on 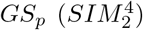 (all genomes were included in the training set) as the baseline. In 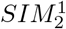 (contaminated genomes excluded in the training set), the F1-score reduction from phylum to species rank was the smallest, with only a 1.07% decrease at the phylum rank and a 17.4% decrease at the species rank. Conversely, 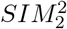 (both core and contaminated genomes were excluded in the training set) exhibited the greatest reduction, with a 19.65% decrease in F1-score at the phylum rank and a 24.48% decrease at the species rank. The observations aligned with our expectations since understanding the sequences’ pattern of the core genomes was essential for the purification of MAGs. Furthermore, we noted a slight decrease of merely 1.07% and 6.81% in the F1-scores for 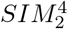 when the lineages were different from phylum to family. Nonetheless, a substantial disparity of 11.84% for genus and 17.14% for species was observed. This finding indicates that Deepurify exhibited greater efficacy in removing contamination when it occurs at higher ranks, irrespective of their inclusion in the training set. In contrast, addressing contamination at lower taxonomic ranks proved to be more challenging due to the increased presence of homologous sequences.

### The impact of homologous sequences and contig length on MAG decontamination

For a simulated MAG, we defined the contigs as derived from homologous sequences if they could be aligned to both core and contaminated genomes (**Methods**). In the test set of *SIM*_1_, we identified contigs from homologous sequences at various taxonomic ranks: 142 at phylum, 832 at class, 3,015 at order, 4,429 at family, 8,048 at genus, and 17,169 at species. The number of contigs from homologous sequences increased from phylum to species, which could explain the reason for the performance declination of MAGpurify, MDMcleaner, and Deepurify if contamination derived from the LCAs of genomes at low taxonomic ranks.

Furthermore, we categorized the contigs based on their lengths (intervals of 1,000 bps) to assess the influence of contigs’ length on the performance of Deepurify. Deepurify showed better performance on long contigs compared to short ones (**Supplementary Figure** 8, **Supplementary Table** 7) probably because long contigs could provide more information on the genomic context of their source genomes.

### Deepurify improves the qualities of MAGs from different contig binning tools

We applied MAGpurify, MDMcleaner, and Deepurify to the MAGs generated by MaxBin, MetaBAT2, VAMB, and CONCOCT to examine if they could increase the number of medium- and high-quality MAGs. The contigs of CAMI I and human gut metagenomic sequencing (*S*1) were downloaded from our previous study [25] (**Methods**). We used two criteria to evaluate the performance of MAG decontamination: 1. the increased number of medium- (*INM*_*mq*_) and high-quality MAGs (*INM*_*hq*_); 2. the improved quality score (*IQS* ), which measures the overall MAG quality improvement (**Methods**). Deepurify consistently outperformed the other two tools in nearly all datasets and contig binning methods (Figure 5, **Supplementary Table** 8). On average, across all datasets and binning methods, Deepurify exhibited 2.87-fold (1.33-fold) and 5.15-fold (4.16-fold) higher mean value for the *INM*_*hq*_ and *INM*_*mq*_ compared to MAGpurify (MDMcleaner). Interestingly, the *INM*_*hq*_ and *INM*_*mq*_ values of Deepurify for all binning tools were commonly positive except for VAMB on the high-complexity community in CAMI I (VAMB does not work well on a single sample) and the values for MAGpurify and MDMclean were more frequently to be observed negatively. In Figure 6, we depict the completeness and contamination rates of MAGs, before and after purification with MAGpurify, MDMcleaner, and Deepurify, using data from CAMI I and *S*1. We also employ a generalized additive model to create a smooth curve, effectively capturing the contamination trends within these MAG datasets. It was observed that Deepurify consistently outperformed the others by having the smallest areas under the curve.

**Figure 5:**
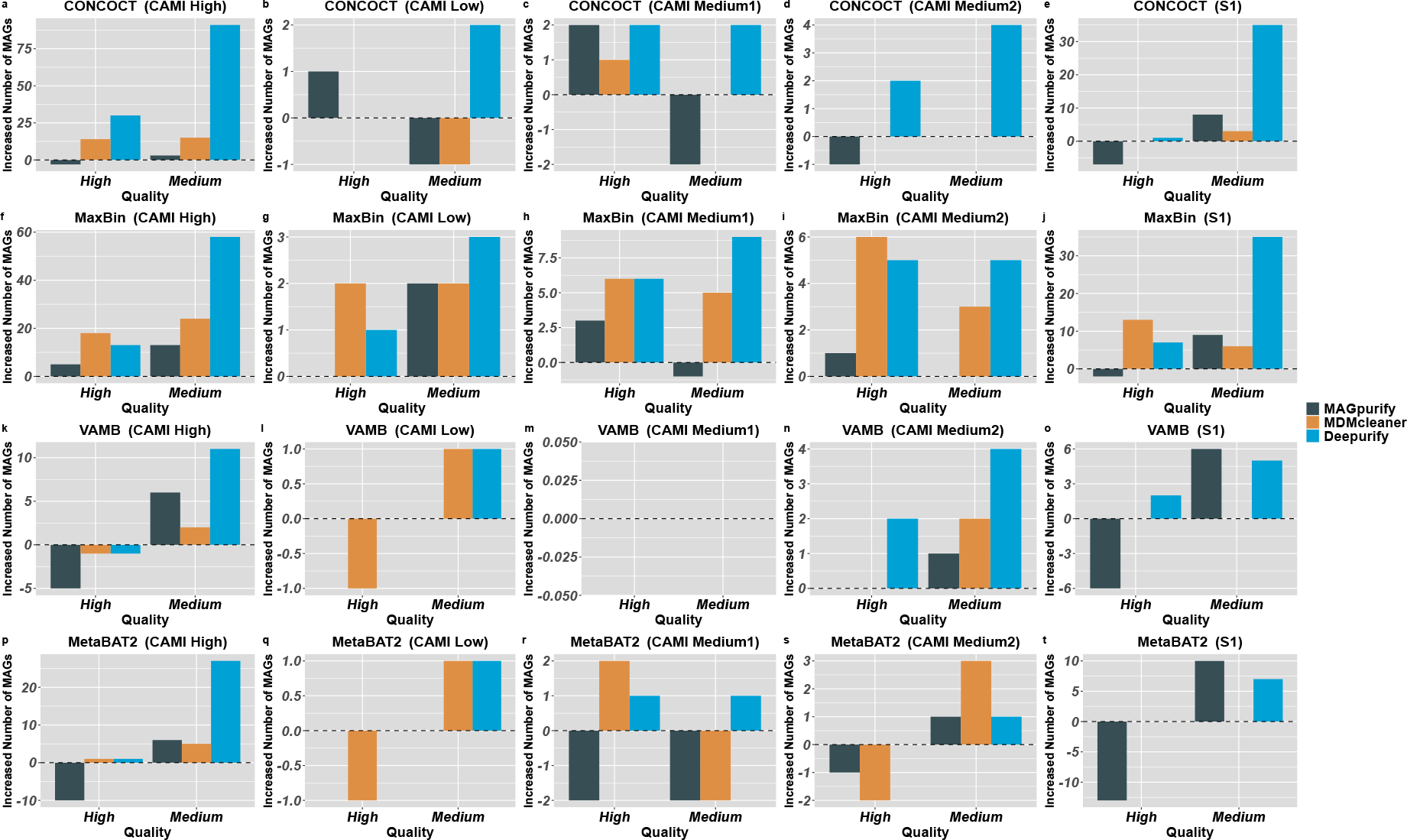
The increased number of MAGs (INM) for CAMI I and *S*1 datasets with different binning methods (CONCOCT, MaxBin, VAMB, MetaBAT2) for MAGpurify, MDMcleaner, and Deepurify.

**Figure 6:**
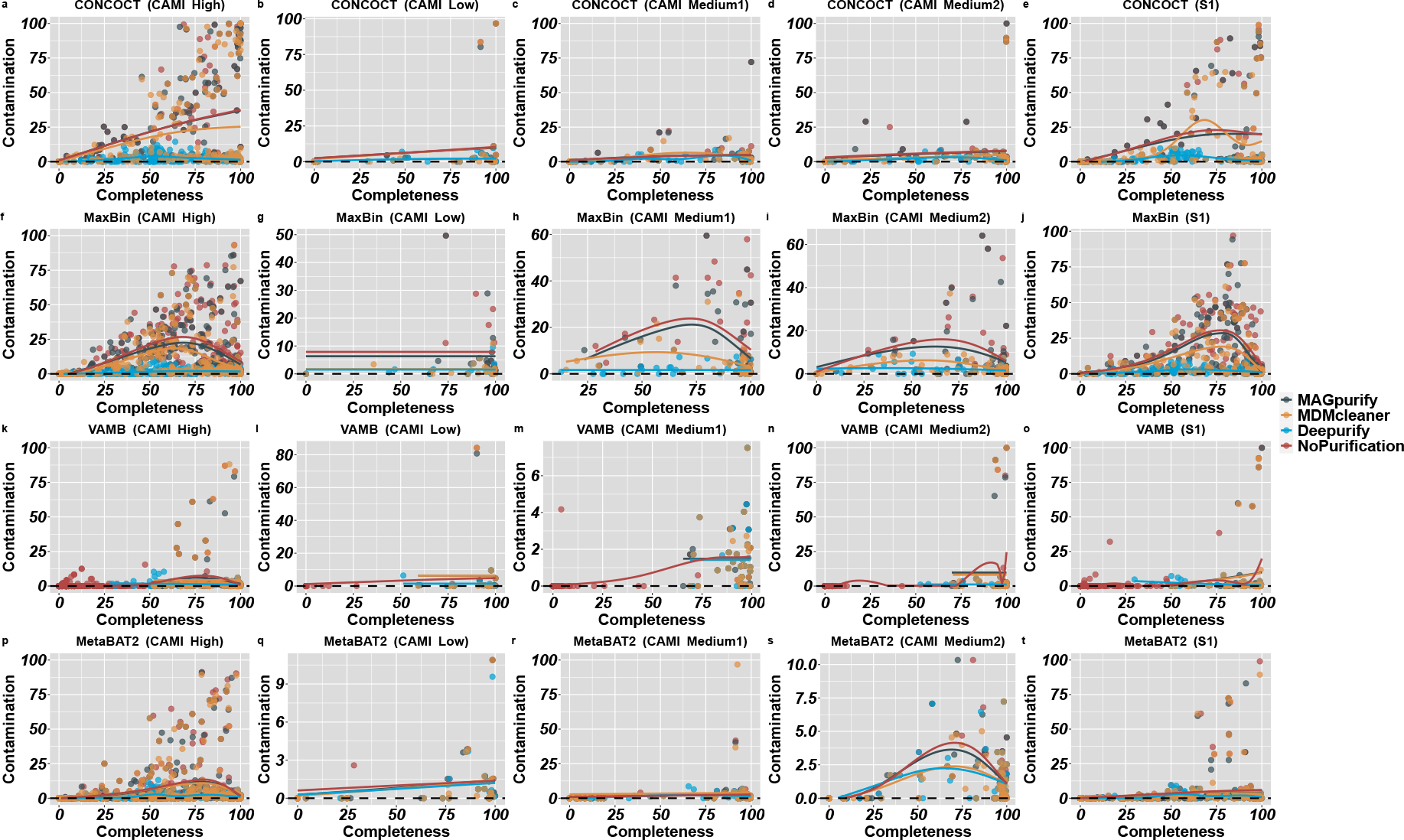
The correlation between the completeness and contamination levels of MAGs both before and after purification using MAGpurify (grey), MDMcleaner (orange), and Deepurify (blue) in the CAMI I and *S*1 datasets. These datasets were initially binned using CONCOCT, MetaBAT2, VAMB, and MaxBin. A Generalized Additive Model (GAM) was applied to construct a smooth curve that represents the contamination trends exhibited by MAGs in these instances. These plots serve to illustrate the superior purification performance of Deepurify when used on MAGs with high contamination levels.

Deepurify demonstrated remarkable performance superiority over MAGpurify (IQS: 29.21-fold on average for all cases) and MDMcleaner (IQS: 1.82-fold on average for all cases), especially on the binning results of CONCOCT, VAMB, and MetaBAT2 (Figure 7, **Supplementary Table** 9). These observations suggested that Deepurify was more effective in improving contig binning performance than other tools as many low-quality MAGs were able to be upgraded to medium- or high-quality MAGs.

**Figure 7:**
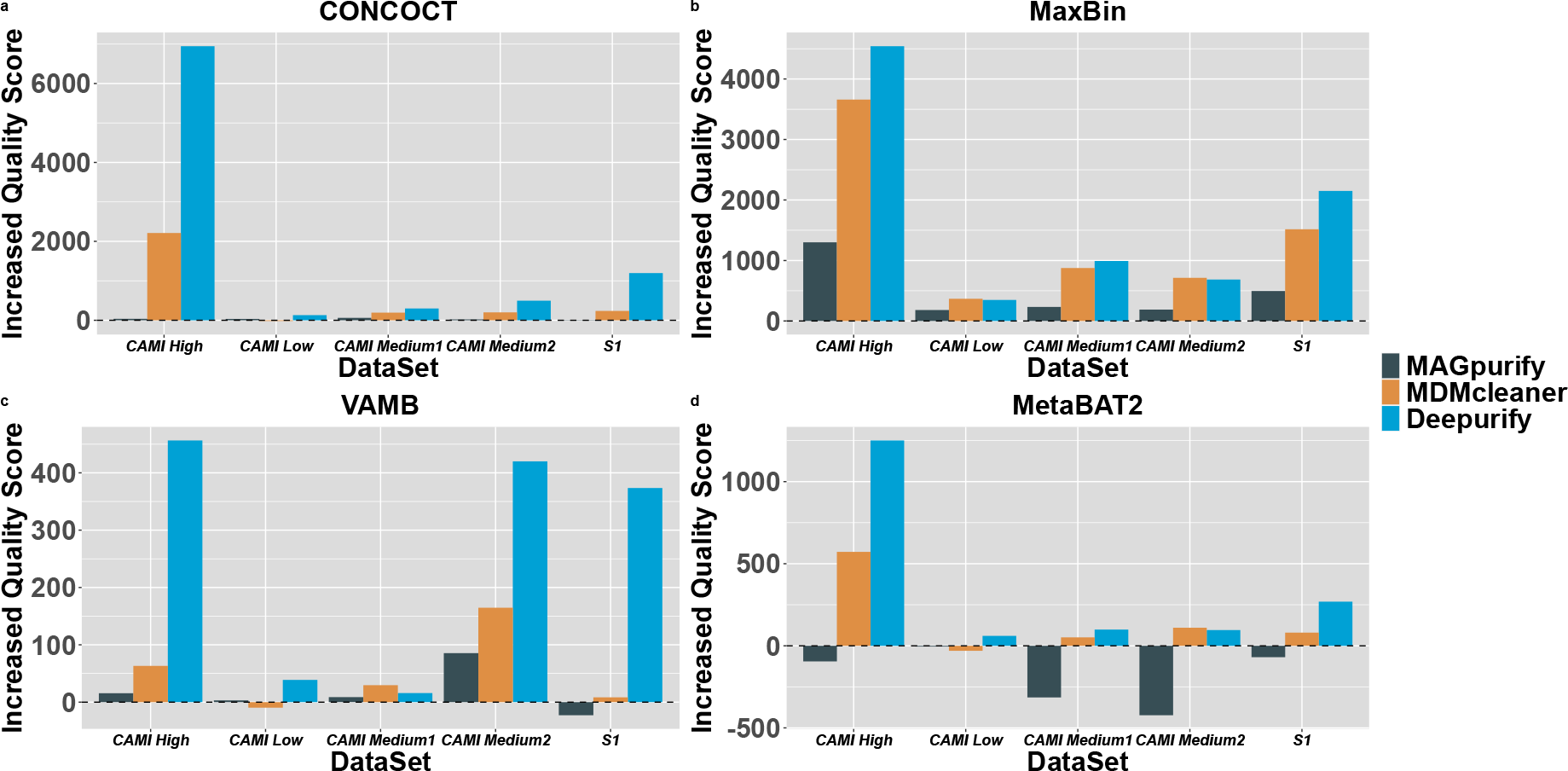
The increased quality scores (IQS) for the CAMI I and *S*1 datasets binned with MaxBin, CONCOCT, VAMB, and MetaBAT2 reveal that Deepurify’s IQS is substantially higher than that of MAGpurify and MDMcleaner in almost all cases.

### Deepurify outperforms other purification tools on real-world data

We further applied Deepurify to the human gut metagenomic sequencing data from 290 IBS-D patients and 89 healthy controls [26]. The sequencing data were assembled by metaSPAdes [5] followed by contig binning using MetaBAT2 (**Methods**), which generated 4,887 high-quality and 5,943 medium-quality MAGs. We selected 713 MAGs with high contamination (completeness ≥ 50% and contamination ≥ 25%) to evaluate the efficacy of Deepurify on MAG decontamination. Our examination revealed that MAGpurify and MDMcleaner could enhance the quality of only a small fraction of these highly contaminated MAGs. Specifically, MAGpurify improved 1.4% of them to high- and medium-quality MAGs (*INM*_*hq*_ = 1, *INM*_*mq*_ = 9), while MDMcleaner improved 0.7% of them to high- and medium-quality MAGs (*INM*_*hq*_ = 1, *INM*_*mq*_ = 4). Deepurify demonstrated a remarkable ability for MAG decontamination, as it was able to rescue a significant proportion of these MAGs, 70.12% of them (*INM*_*hq*_ = 3, *INM*_*mq*_ = 497). Deepurify demonstrated a significantly elevated IQS at 248994.46, surpassing both MAGpurify, which has an IQS of 17772.28, and MDMcleaner, with an IQS of 14466.47. The contamination rates of these MAGs were mostly reduced to below 10% after undergoing MAG decontamination using Deepurify whereas the values obtained from the other tools were considerably higher (Figure 8 **a**).

**Figure 8:**
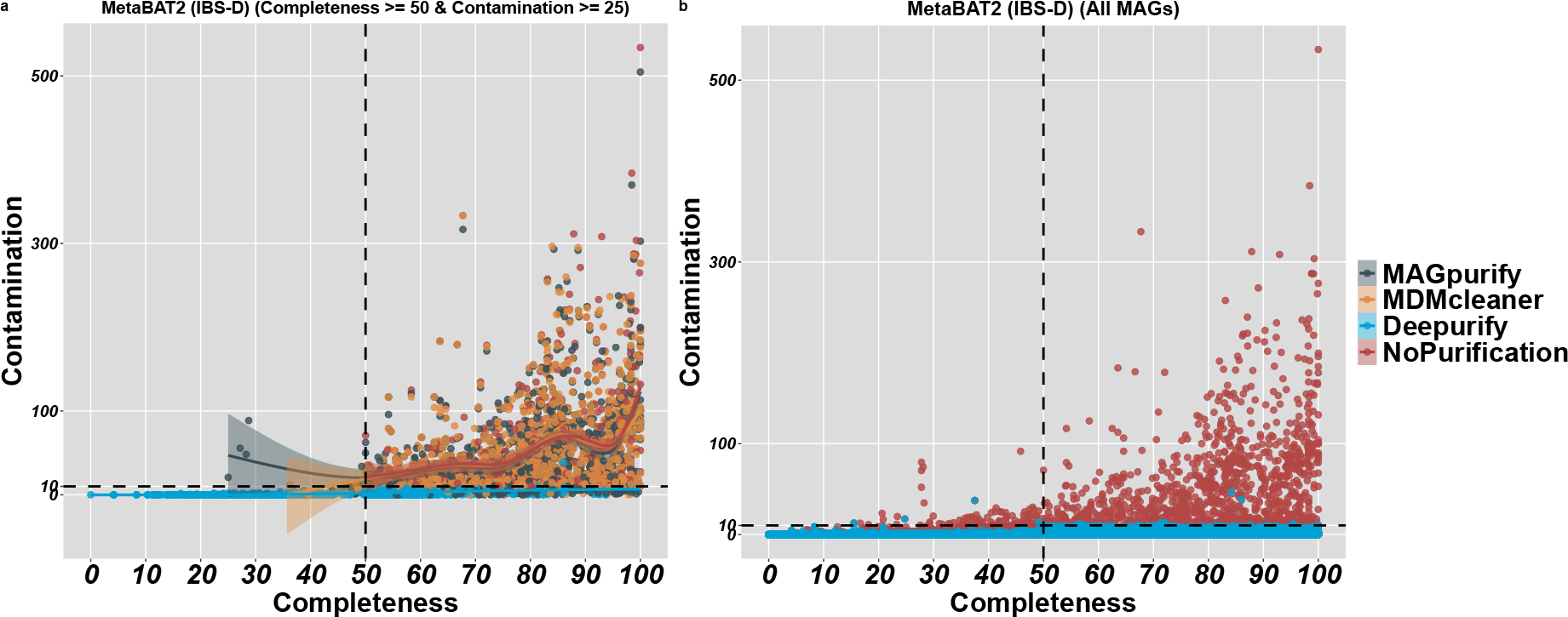
The correlation between completeness and contamination of MAGs before and after purification. In (**a**), we employed MAGpurify (grey), MDMcleaner (orange), and Deepurify (blue) to filter the contamination of MAGs with completeness greater than 50% and contamination exceeding 25%. A Generalized Additive Model (GAM) was applied to construct a smooth curve that effectively captured the contamination trends exhibited by MAGs in these instances. In (**b**), Deepurify (blue) was utilized for all MAGs within the IBS-D cohort. Notably, Deepurify exhibits the capacity to rescue a significant proportion of MAGs with high contamination rates (*>* 10%).

### Deepurify identified novel IBS-D association signals

We examined all high-quality (4,931) and medium-quality (6,539) MAGs obtained from the IBS-D cohort after Deepurify decontamination to identify novel association signals. We utilized GTDB-TK [**29**](**Methods**) to annotate these MAGs both before and after Deepurify’s purification process. Upon comparing the MAG annotation results, we identified five new species (**Supplementary Table** 5) and one new genus (**Supplementary Table** 6). We performed an association analysis of IBS-D on the 678 MAGs, which were initially categorized as low-quality but were reclassified as medium- or high-quality after decontamination (**Methods**). This analysis identified several suggestive signals (P-value *<* 0.05) including one novel species (s Collinsella sp900541055), and two confirmed species (Alistipes[30]and Ruminococcus gnavus [31, 32]) that were known to be associated with IBS-D. Lastly, we showed the completeness and contamination rates for all MAGs in the IBS-D cohort before and after purification by Deepurify in Figure 8 **b**. This plot demonstrated Deepurify’s remarkable ability to purify contaminated contigs in MAGs.

## Discussion

Utilizing genome assembly with short-read metagenomic sequencing data has become a prevalent method to decipher microbial compositions in complex environments. However, each assembled metagenomic contig only partially represents a microbial genome. It is therefore crucial to perform contig binning to obtain contig sets with similar genomic characteristics and abundances, which then represent MAGs that originate from the same microbe. As was highlighted in a recent paper [15], MAG contamination is a significant stumbling block during contig binning on single sample assembly. Decontamination tools, such as MAGpurify and MDMcleaner, have been developed to address the challenge of eliminating contaminated contigs from MAGs. Nonetheless, these tools demonstrate several limitations. Most notably, they are ineffective in distinguishing contigs from each other if their core and contaminated genomes belong to the same family or genus. Furthermore, these tools are unable to process contigs whose source genomes are absent from their built-in databases. And thirdly, they mainly focus on genes, with genomic variations such as gene order and genome rearrangements being left out of consideration.

To address these limitations, we developed Deepurify, a novel tool that uses deep language models to learn the relationship between microbial genomes and their taxonomic lineages. Deepurify models all nucleotides in sequences and can learn local genomic alternations between adjacent species in the training set. This approach allows Deepurify to handle contigs without known genes. Deepurify has a superior decontamination capacity, particularly if the contigs share a high proportion of homologous sequences (**Supplementary Note** 4). It also outperformed the existing tools if the source genomes of contigs were not included in the training dataset. Deepurify could significantly speed up the MAG decontamination procedure with GPU acceleration (**Supplementary Note** 2), which allows scaling of decontamination to large numbers of MAGs. The primary runtime bottleneck for Deepify lies in the duration required for running CheckM, which is nearly twice as long as inferring lineages for the contigs within MAGs. The efficiency of Deepurify’s execution could be significantly improved with a method to expedite the CheckM runtime.

Deepurify adopts a unique approach to optimize the utilization of contigs within a MAG. Instead of adopting the common practice of directly discarding contaminated contigs, Deepurify constructs a MAG-separated tree for filtering. This innovative strategy proves especially advantageous in scenarios where MAGs exhibit a substantial degree of contamination, typically exceeding a contamination rate of 100%. Deepurify has the ability to resolve a highly contaminated MAG into two separate MAGs, typically falling within the high- or medium-quality range. On occasion, it may yield three or more MAGs that hold potential for further utilization.

Our experiments demonstrated the remarkable efficacy of Deepurify in decontaminating MAGs from short-read assembly. We hold a strong belief that its applicability extends to contigs derived from long-read assemblies, accompanied by two distinct advantages: Firstly, contigs derived from long-read assemblies are significantly longer than those from short-read assemblies. It offers Deepurify a substantially enriched sequence context, thereby enhancing its capacity for decontamination. Secondly, single-base substitutions and indel errors are frequently observed in long-read assemblies [33], which we placed emphasis on during the development of Deepurify’s training procedure (**Supplementary Note** 6). It is worth mentioning that contemporary decontamination tools do not typically consider sequence noise.

On the other hand, it is important to note that Deepurify cannot deal with overly large misassemblies in contigs (such as chimeric contigs, translocations, etc.). We observed that for some MAGs Deepurify failed to achieve its specified decontamination standard because a designated single-copy gene was detected multiple times. Chimeric contigs may therefore remain a challenge to Deepurify since they could substantially influence the local context of sequences, which may adversely impact the quantification of taxonomic similarity between contigs in a MAG. To mitigate the influence of such misassemblies, we recommend that users apply assembly error correction tools such as metaMIC [34] prior to using Deepurify.

## Methods

### Preparing and processing microbial reference genomes

We downloaded microbial representative genomes and their taxonomic lineages from proGenomes v2.1 database [35] to generate two training sets *GS*_*c*_, *GS*_*p*_ for model training and two simulated testing sets *SIM*_1_ and *SIM*_2_ for evaluating. We excluded the microbial genomes without phylum annotations or if the phyla they belonged to had less than 15 species. For microbes with only phylum and species annotations, all other taxonomic ranks inbetween were annotated as “Unclassified”.

### Training sets construction

After data preprocessing, we generated two training sets for *SIM*_1_ and *SIM*_2_: 1. a complete reference genome training set (*GS*_*c*_) consisting of the genomic sequences from 10,332 species belonging to 37 phyla (**Supplementary Table** 10); 2. a partial reference genome training set (*GS*_*p*_) by randomly selecting 112 species, which come from 12 phyla (**Supplementary Table** 11) in *GS*_*c*_. *GS*_*p*_ was used to evaluate the performance of Deepurify when either core or contaminated genomes were not included in the training set.

During the training stage, we sampled the contig-sized sequences from the genomes in *GS*_*c*_ and *GS*_*p*_. The sequence lengths ranged from 1,000 bps to 8,192 bps, following a pre-defined contig length distribution learned from a real metagenomic assembly exercise (**Supplementary Note** 5). We randomly incorporated into these sequences insertions, deletions, and single nucleotide variants (**Supplementary Note** 6) in order to reduce the impact of sequencing errors and enhance model generalization capabilities.

### **MAG generation for** *SIM*_1_ **and** *SIM*_2_

We simulated chimeric MAGs for use in *SIM*_1_ and *SIM*_2_ sets for evaluation: 1. For *SIM*_1_, all source genomes of contigs were included in *GS*_*c*_; 2. For *SIM*_2_, some source genomes of contigs might be absent from *GS*_*p*_. For *SIM*_1_, we randomly selected the genomes of two distinct lineages (*SP*_1_ and *SP*_2_) in *GS*_*c*_ and simulated 200 contigs from them with lengths between 1,000 bps and 8,192 bps and with varying proportions of contigs from *SP*_2_ (5%, 10%, 15%, and 20%) for each MAG. The lineage LCAs of *SP*_1_ and *SP*_2_ were traversed from kingdom to genus (lineages differ starting from phylum to species). We generated 50 MAGs for each mixture proportion and on each taxonomic rank of LCA. For *SIM*_2_, we followed a similar chimeric MAG simulation procedure as we did for *SIM*_1_, the only difference being that *SP*_1_ and *SP*_2_ may be extracted from either *GS*_*p*_ or from *GS*_*c*_ − *GS*_*p*_. There are four permuted scenarios for *SIM*_2_: 1. both core and contaminated genomes are included in the 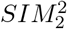. only core genomes are included in the 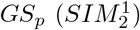 3. only contaminated genomes are included in the 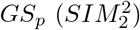 4. neither core nor contaminated genomes are included in the 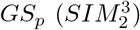.

### Generate MAGs from contig binning tools

We downloaded the metagenomic sequencing datasets from CAMI I with low, medium (two datasets), and high complexity and from a human stool sample (*S*_1_) [25]. The contigs of these datasets were assembled by metaSPAdes with default parameters. Contigs were grouped as MAG using VAMB (contig length *>* 1kbps), CONCOCT (contig length *>* 1kbps), MaxBin (contig length *>* 1kbps), and MetaBAT2 (contig length *>* 1.5kbps). We only kept MAGs from VAMB with a completeness of at least 50% to exclude the MAGs with few contigs (e.g. *<* 3 contigs).

### MAG quality definitions

MAGs are typically classified into distinct quality categories based on their degrees of completeness and contamination. High-quality MAGs are defined by completeness levels equal to or exceeding 90% and contamination levels at or below 5%. Medium-quality MAGs are characterized by completeness levels of 50% or higher, with contamination levels below or equal to 10%. MAGs failing to meet the high or medium-quality criteria are categorized as low-quality.

### IBS-D real-world validation study

We applied metaSPAdes with default parameters to assemble short-read metagenomic sequencing data from 290 IBS-D patients and 89 healthy controls. The contigs longer than 1.5kb were grouped into MAGs by MetaBAT2. We evaluated MAGpurify, MDMcleaner, and Deepurify on the MAGs with completeness ≥ 50% and contamination ≥ 25%.

### Microbial taxonomic annotation

We used GTDB-Tk [29] to annotate and allocate MAGs to the taxonomic tree. A MAG would be annotated as a particular species (*g*_*ref*_ represents its genome) if 1. its average nucleotide identity with *g*_*ref*_ is no less than 95% and 2. its alignment fraction against *g*_*ref*_ is no less than 65%.

### Identification of homologous sequences

We conducted BLASTN alignments between the core contigs and the contaminated genomes, as well as between the contaminated contigs and the core genomes. Contigs with an *E*-value less than 1*e*^*−*6^ were considered aligned and would be categorized as sequences derived from homologous sequences.

### Metrics for performance evaluation

For simulated chimeric MAGs, we applied a balanced macro F1-score to evaluate the performance of MAG decontamination to mitigate the influence of unbalanced numbers of contigs from *SP*_1_ and *SP*_2_. For the binned MAGs that were generated from CAMI I, *S*_1_ and the IBS-D cohort, we adopted two criteria to evaluate the improvement of MAG qualities: 1. total increased number of high-(completeness ≥ 90% and contamination ≤ 5%) and medium-quality (completeness ≥ 50% and contamination ≤ 10%) MAGs; 2. increased quality score (IQS)

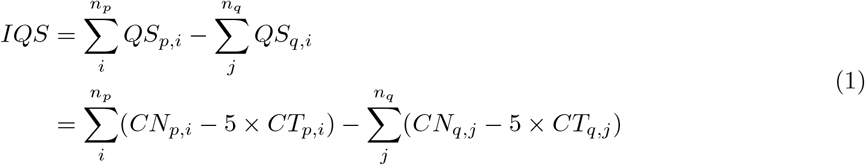

where *n*_*p*_ and *n*_*q*_ denote the total number of high- and medium-quality MAGs after and before MAG decontamination, respectively. *CN*_*p,i*_ (*CN*_*q,j*_) and *CT*_*p,i*_ (*CT*_*q,j*_) are the completeness and contamination values of the MAGs after (before) MAG decontamination.

### Architecture of Deepurify

#### Genomic sequence and taxonomic lineage encoders

Deepurify utilizes GseqFormer and LSTM to encode genomic sequences and taxonomic lineages into 1024-dimensional space. The fundamental architecture of Deepurify is illustrated in Figure 1 **b**.

#### GseqFormer: Genomic sequence encoder

The genomic sequences were represented as a unified embedded matrix by concatenating the sequence representations with one-hot, 3-mers, and 4-mers (**Supplementary Note** 7). We developed GseqFormer to encode the sequence-embedded matrix in high dimensional space. It was built on the structure of UniFormer [36], which takes advantage of transformer and convolutional neural networks (CNNs). We substituted the attention module of UniFormer with a new gated self-attention module, which was modified from Evoformer [37] (**Supplementary Note** 8). Because UniFormer has a limitation in modeling long sequences (1,000 bps), we adopted EfficientNet [38] to compress the input sequences into 512 tokens. This strategy enables the maximum lengths of input sequences up to 8,192 bps. Additionally, we incorporated a variety of *tricks* [39, 40, 41] into EfficientNet for efficient training and improved model robustness (**Supplementary Note** 9).

#### LSTM: Taxonomic lineage encoder

The taxonomic lineage 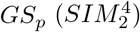of a sequence (*s*_*i*_) at species rank was considered as a sentence that concatenates taxon 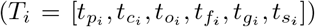 at different taxonomic ranks (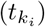, spanning from phylum to species. The taxonomic sentence would be encoded by a 5-layer LSTM model.

### Deepurify training procedure

#### Contrastive training

For a sequence *s*_*i*_, we represented its normalized encoded vector as 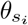 and the normalized encoded vector of its taxonomic lineage at the species rank as 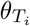.The prefix of *T*_*i*_ before *k*_*i*_ rank denotes as 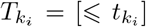We leveraged contrastive training to enable Deepurify to discriminate true (positive label, 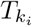) and multiple fake (negative labels, 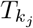) taxonomic lineages for a given sequence *s*_*i*_. During training, we randomly selected *k*_*i*_ and created fake taxonomic lineages 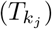 from the taxonomic tree for contrastive, making sure they were distinct from 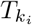 (**Supplementary Note** 10).

We applied four loss functions in contrastive training, including 1. sequence-taxonomy (ST) loss, lineage-phyla (LP) loss, 3. indel loss, and 4. phyla-rank (PR) loss. Deepurify’s primary objective is to optimize ST loss, which aims to make 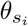 have higher cosine similarity with 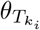 than with 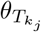. The ST loss (*L*_*ST*_ ) is defined as:

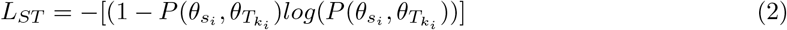

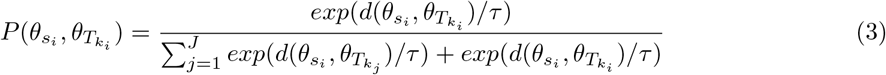

where 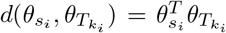), *τ* is a learnable parameter, *J* is the number of negative labels used in contrastive training. The numbers of species are different across phyla, which leads to unbalanced sequences in the training set. We applied an oversampling strategy (**Supplementary Note** 11) and the focal loss [42] to mitigate this problem.

The goal of LP loss (*L*_*LP*_ ) is to establish a taxonomic encoder to minimize the distance between 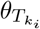 and the phylum that *s*_*i*_ belongs to 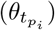.

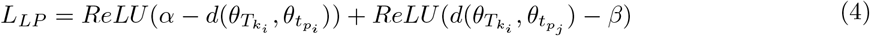

where *α* and *β* are between 0 and 1, which control the cosine similarities between 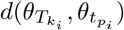 and 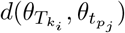.

The aim of indel loss was to enable GseqFormer to accept the sequences with insertions and infer masked sequences.

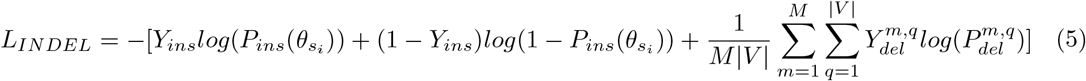

where 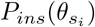 is the predicted probability of *s*_*i*_ including insertions, and *Y*_*ins*_ = 1 indicates *s*_*i*_ including insertions. *M* is the number of masked positions in *s*_*i*_ and each position has six candidate values (*V* = {*A, T, C, G, N, padding*}).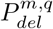 is the predicted probability of *m*-th masked position equals to 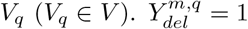 if the *m*-th masked position is *V*_*q*_.

PR loss was used to examine the taxonomic inference of Deepurify on phylum rank.

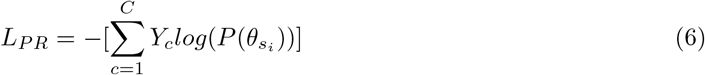

where *C* is the number of phyla in the taxonomic tree, *Y*_*c*_ = 1 if *s*_*i*_ belongs to the phylum *c*, 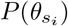the predicted probability of *s*_*i*_ belongs to the phylum *c*.

Therefore, the final training loss function of Deepurify is defined as follows:

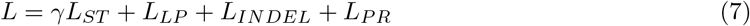

We set *γ* = 2 in our experiments to emphasize the importance of *L*_*ST*_ in Deepurify training. The settings of other hyper-parameters were similar to UniFormer [36] (**Supplementary Note** 12).

### Deepurify MAG decontamination procedure

#### Quantifying sequence taxonomic similarity

Deepurify utilized GseqFormer to encode genomic sequences and then to quantify their taxonomic similarities. This is achieved by identifying the taxonomic lineage from the taxonomic tree that exhibited the highest similarity with the genomic sequences (Figure 2 **a**). The degree of similarity between the sequences is positively correlated with the similarity of their predicted taxonomic lineages. For sequence *s*_*i*_, GseqFormer would calculate the 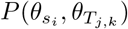for every taxon *j* at taxonomic rank *k*, where *T*_*j,k*_ = [ < *t*_*k*_,*j*], *n* is the total number of taxa in rank *k*. We then selected the three candidate taxa with the highest values. The calculation of 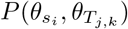 is similar to Eq.(3). For rank *k* + 1, Deepurify would only search for the nodes, whose parents have been selected in rank *k*. This top-*k* searching strategy would result in a number of paths, *ω*, from the root to the species rank (*T*_*j*_, *j* = 1舰*ω*). We then calculate 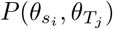 to select the best path.

#### Detecting contaminated contigs in simulated MAGs

On simulated data, a contig with low taxonomic similarity to others in a MAG is more likely to be contaminated. Consequently, contigs were classified as contaminants if their predicted lineages differed from the predominant ones (Figure 2 **b**). We collected the predicted taxonomic lineages of contigs in a MAG and implemented an approach to determine the predominant one. The *Score*_*j,k*_ was calculated for taxon *j* at rank *k*,

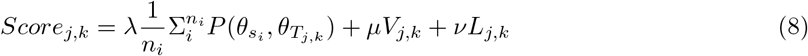

where *n*_*i*_ is the number of contigs that have predicted annotation *j* at rank *k. V*_*j,k*_ and *L*_*j,k*_ denote the proportions of contigs and their total length in a MAG with the taxonomic lineage of *T*_*j,k*_, respectively. We would select *T*_*j,k*_ with the highest value as the predominant lineage in the MAG at rank *k*. The selection would be performed for each rank, where the selected predominant lineage at rank *k* + 1 should be the offspring of the one at rank *k*. At rank *k*, the contigs were identified as contaminants if their predicted lineages were different from the predominant ones.

#### Optimizing contig utilization in MAGs

On real data, Deepurify divides the contigs from a MAG to maximize the number of medium- and high-quality MAGs using the MAG-separated tree. The MAG-separated tree is constructed based on the predicted taxonomic lineage for the contigs in a MAG (Figure 2 **c**). Each node includes the contigs with the same annotation at rank *k*. We collected single-copy genes (SCGs) from the databases of SolidBin [43] and bacteria and archaea domains in CheckM [28]. We used Prodigal [44] to predict genes on contigs and aligned them with SCGs by HMMER (http://hmmer.org). We removed contigs to eliminate duplicated SCGs within each node (**Supplementary Note** 13). This procedure may result in multiple candidate contig divisions for a node. To enhance computational efficiency, Deepurify discarded the divisions if 1. more than 45% of the original SCGs were removed and 2. the total lengths of involved contigs were less than 550kb (**Supplementary Note** 14). We applied CheckM to each division and selected the best one to represent a node based on quality score (*QS*). Its quality (high-, medium- or low-quality) was also annotated by CheckM. Deepurify utilized post-order traversal to traverse the MAG-separated tree to maximize the total number of medium- and high-quality MAGs (Figure 2 **d**, **Supplementary Note** 15).

## Data availability

The microbial representative genomes and their associated taxonomic lineages were downloaded from the proGenomes v2.1 database. The *SIM*_1_ was uploaded to https://zenodo.org/record/8343498. The *SIM*_2_ was uploaded to https://zenodo.org/record/8343506. The CAMI I short-reads were downloaded from ‘1st CAMI Challenge Dataset 1 CAMI low’, ‘1st CAMI Challenge Dataset 2 CAMI medium’ and ‘1st CAMI Challenge Dataset 3 CAMI high’ with the following link: https://data.cami-challenge.org

/participate/. The Illumina short-reads, 10x linked-reads, and long-reads of *S*1 data were downloaded from NCBI SRA accessions SRR19505636. The fecal metagenomic sequencing reads of the IBS-D cohort were downloaded from China National GeneBank (CNGB) with accession number CNPO000334.

## Code availability

The source code used in the manuscript is freely available under an MIT license at https://github.com/zoubohao/Deepurify_Project. The versions of the software used in the study were provided in the **Supplementary Note** 16.

## Authors’ contributions

LZ conceived the study. BHZ designed and implemented the Deepurify algorithms. LZ and BHZ conceived the experiments. BHZ, YD, and ZMZ conducted the experiments. BHZ and JJW analyzed the results. BHZ drew and analyzed the plots. BHZ and LZ wrote the manuscript. KCC and SS contributed computational resources.

