## Supplementary File for "Deepurify: a multi-modal deep language model to remove contamination from metagenome-assembled genomes"

### Supplementary Notes

---

#### 1 An analysis of MAG abundances with both high and low levels of contamination

We selected all MAGs with high levels of contamination (Completeness $\geq$ 50% and Contamination $>$ 10%) and randomly selected 1,000 MAGs with low levels of contamination (Completeness $\geq$ 50% and Contamination $\leq$ 10%) from IBS-D cohort. MAG abundance estimation was conducted through the mapping of reads to MAG contigs. The Reads Per Kilobase per Million mapped read (RPKM) of these MAGs were calculated.

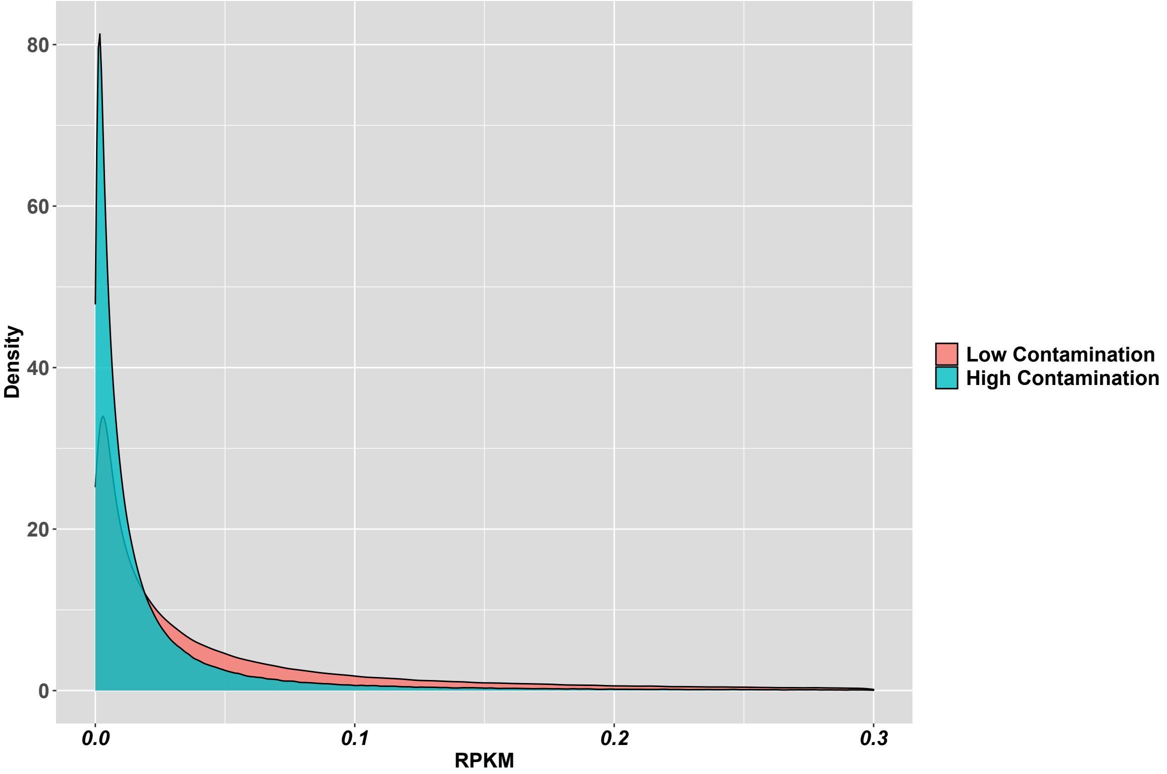


Supplementary Figure 1: The density distribution of RPKM for MAGs with high and low contamination

The distribution of RPKM values for MAGs with high and low contamination exhibited significant overlap, as depicted in **Supplementary Figure** 1. This suggested that some MAGs with high abundances may not be suitable for further analysis due to contamination, potentially leading to the exclusion of MAGs that exhibited significant associations with IBS-D.

#### 2 Comparison of running time of MDMcleaner and Deepurify

We conducted a runtime comparison between Deepurify and MDMcleaner using the CAMI I and $S1$ datasets. We excluded MAGpurify from this comparison due to its inferior decontamination performance in our study. Both MDMcleaner and Deepurify utilized four threads for MAG purification (inference). Deepurify, in particular, efficiently utilized two threads per GPU, making use of the computational power of two NVIDIA GTX 3090 24GB GPUs.

During the evaluation stage, MDMcleaner employed a single CheckM process with seventy-five threads for evaluating MAG quality. In contrast, Deepurify's evaluation stage applied three parallel CheckM processes, each utilizing twenty-five threads, to evaluate all MAG nodes' quality in the MAG-separated tree. We recorded the inference and evaluation times for both MDMcleaner and Deepurify.

| ***Purification Tool*** | MDMcleaner | | | Deepurify | | |
|---|---|---|---|---|---|---|
| Dataset | Inference | Evaluation | Total | Inference | Evaluation | Total |
| CAMI I High (CONCOCT) | 10452m | **61m** | 10513m | **491m** | 1004m | **1495m** |
| CAMI I Medium 1 (CONCOCT) | 878m | **25m** | 903m | **49m** | 88m | **137m** |
| CAMI I Medium 2 (CONCOCT) | 909m | **29m** | 938m | **59m** | 175m | **234m** |
| CAMI I Low (CONCOCT) | 957m | **16m** | 973m | **29m** | 116m | **146m** |
| $S_1$ (CONCOCT) | 4477m | **40m** | 4517m | **123m** | 296m | **420m** |
| CAMI I High (MaxBin) | 10889m | **46m** | 10935m | **282m** | 673m | **955m** |
| CAMI I Medium 1 (MaxBin) | 1632m | **21m** | 1653m | **44m** | 135m | **179m** |
| CAMI I Medium 2 (MaxBin) | 2129m | **25m** | 2154m | **53m** | 178m | **231m** |
| CAMI I Low (MaxBin) | 817m | **15m** | 832m | **19m** | 82m | **101m** |
| $S_1$ (MaxBin) | 8027m | **41m** | 8068m | **118m** | 300m | **418m** |
| CAMI I High (VAMB) | 4566m | **34m** | 4600m | **117m** | 443m | **560m** |
| CAMI I Medium 1 (VAMB) | 867m | **20m** | 887m | **22m** | 102m | **124m** |
| CAMI I Medium 2 (VAMB) | 966m | **19m** | 985m | **27m** | 126m | **153m** |
| CAMI I Low (VAMB) | 409m | **14m** | 423m | **17m** | 79m | **96m** |
| $S_1$ (VAMB) | 1332m | **24m** | 1356m | **27m** | 129m | **156m** |
| CAMI I High (MetaBAT2) | 10749m | **73m** | 10822m | **282m** | 673m | **955m** |
| CAMI I Medium 1 (MetaBAT2) | 1255m | **14m** | 1269m | **33m** | 122m | **156m** |
| CAMI I Medium 2 (MetaBAT2) | 1536m | **24m** | 1560m | **39m** | 162m | **201m** |
| CAMI I Low (MetaBAT2) | 617m | **16m** | 633m | **14m** | 71m | **86m** |
| $S_1$ (MetaBAT2) | 4730m | **37m** | 4767m | **70m** | 213m | **283m** |

Supplementary Table 1: The running time for MDMcleaner and Deepurify in the five datasets. The "m" represents the minute. Bold black fonts indicate a shorter time.

**Supplementary Table** 1 presents the elapsed times for inference and evaluation for both MDMcleaner and Deepurify. Deepurify demonstrated significantly faster inference times, being 35.61 times faster than MDMcleaner, thanks to GPU acceleration. However, its evaluation process was 8.69 times slower because it assessed the quality of each MAG node in a MAG-separated tree. In terms of total time required, Deepurify exhibited an 8.82-fold increase in efficiency compared to MDMcleaner.

#### 3 An analysis of the annotation of contigs in highly contaminated MAGs

We selected 1110 highly contaminated MAGs (Completeness $\geq$ 50% and Contamination $>$ 10%) from the IBS-D cohort, which were assembled and binned using metaSPAdes and MetaBAT2. We utilized Kraken2 [45] with a standard database to annotate the contigs in these MAGs.

We identified the taxon with the highest number of contigs as the predominant taxon, while the remaining taxa were considered contaminated at each taxonomic rank (from phylum to species). The rates of contamination at a taxonomic rank were assessed by calculating the ratio of contigs belonging to contaminated taxa to the total number of annotated contigs.

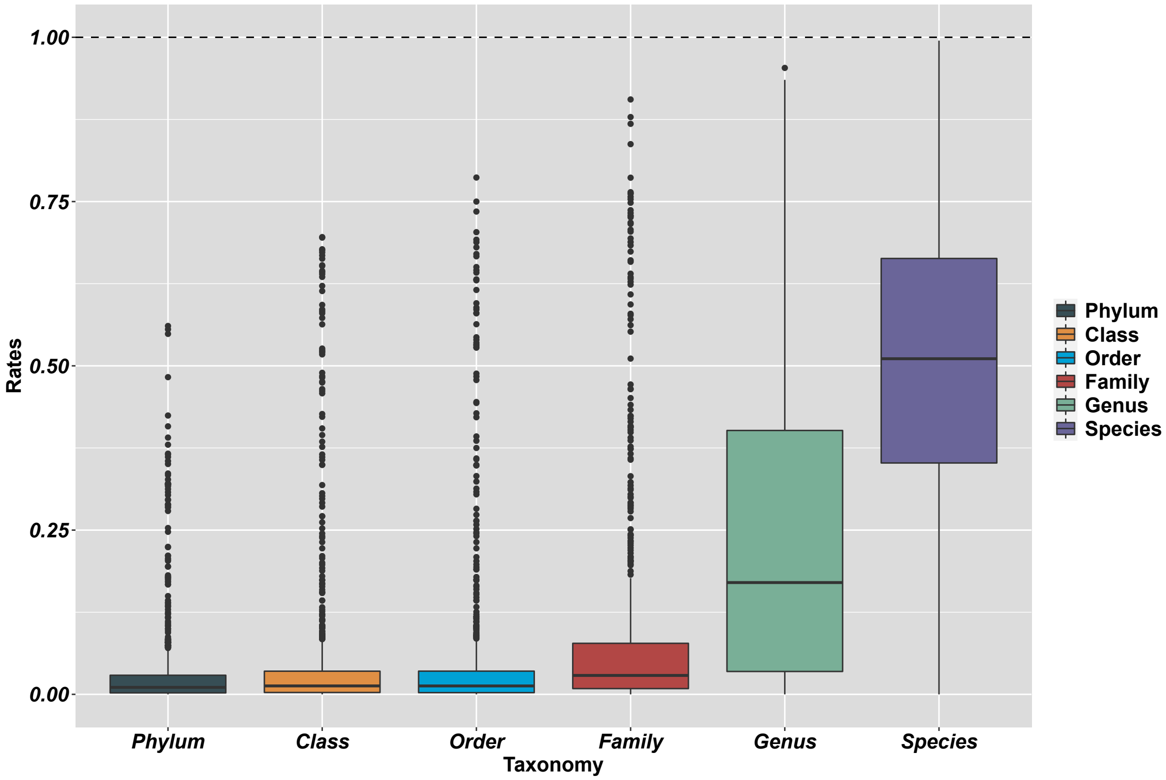


Supplementary Figure 2: The estimated contamination rates at six taxonomic ranks.

**Supplementary Figure** 2 revealed that at higher taxonomic ranks, such as phylum, class, and order, contamination mostly did not occur, but contamination became concentrated at lower taxonomic levels, including family, genus, and species.

#### 4 The averaged precision and recall for MAGpurify, MDMcleaner, and Deepurify on $SIM_1$

We assessed precision and recall for three purification methods: MAGpurify, MDMcleaner, and Deepurify, using the $SIM_1$ simulated testing set. Contaminated contigs were treated as positive labels, while core contigs served as negative labels. We calculated balanced precision and recall to account for the label imbalance in the simulated MAG, where core and contaminated contigs had uneven distributions. This involved assigning higher weights to contaminated contigs, adjusted according to the ratio between the number of core contigs and contaminated contigs within a simulated MAG. **Supplementary Table** 2 summarizes the precision and recall values for each purification method.

| | Contamination 5% | | | | | | Contamination 10% | | | | | |
|---|---|---|---|---|---|---|---|---|---|---|---|---|
| | MAGpurify | | MDMcleaner | | Deepurify | | MAGpurify | | MDMcleaner | | Deepurify | |
| Taxonomy | Precision | Recall | Precision | Recall | Precision | Recall | Precision | Recall | Precision | Recall | Precision | Recall |
| Phylum | 0.921 | 0.592 | 0.991 | 0.988 | 0.977 | 1.000 | 0.980 | 0.577 | 0.998 | 0.993 | 0.988 | 0.999 |
| Class | 0.877 | 0.312 | 0.997 | 0.992 | 0.950 | 0.994 | 0.961 | 0.344 | 0.997 | 0.984 | 0.949 | 0.995 |
| Order | 0.895 | 0.256 | 0.998 | 0.936 | 0.899 | 0.996 | 0.890 | 0.271 | 0.997 | 0.961 | 0.899 | 0.981 |
| Family | 0.891 | 0.158 | 0.978 | 0.700 | 0.848 | 0.966 | 0.961 | 0.220 | 0.998 | 0.710 | 0.873 | 0.966 |
| Genus | 0.865 | 0.094 | 0.918 | 0.480 | 0.812 | 0.978 | 0.916 | 0.154 | 0.853 | 0.293 | 0.820 | 0.920 |
| Species | 0.781 | 0.113 | 0.875 | 0.216 | 0.722 | 0.922 | 0.769 | 0.082 | 0.879 | 0.174 | 0.706 | 0.915 |

| | Contamination 15% | | | | | | Contamination 20% | | | | | |
|---|---|---|---|---|---|---|---|---|---|---|---|---|
| | MAGpurify | | MDMcleaner | | Deepurify | | MAGpurify | | MDMcleaner | | Deepurify | |
| Taxonomy | Precision | Recall | Precision | Recall | Precision | Recall | Precision | Recall | Precision | Recall | Precision | Recall |
| Phylum | 0.976 | 0.424 | 0.999 | 0.995 | 0.986 | 0.999 | 0.958 | 0.385 | 0.996 | 0.996 | 0.981 | 0.998 |
| Class | 0.952 | 0.221 | 0.992 | 0.974 | 0.960 | 0.990 | 0.906 | 0.294 | 0.991 | 0.987 | 0.959 | 0.988 |
| Order | 0.881 | 0.226 | 0.998 | 0.952 | 0.880 | 0.978 | 0.955 | 0.206 | 0.998 | 0.960 | 0.905 | 0.981 |
| Family | 0.926 | 0.138 | 0.976 | 0.779 | 0.860 | 0.975 | 0.947 | 0.186 | 0.977 | 0.792 | 0.887 | 0.974 |
| Genus | 0.879 | 0.163 | 0.839 | 0.328 | 0.816 | 0.922 | 0.960 | 0.107 | 0.879 | 0.482 | 0.785 | 0.920 |
| Species | 0.851 | 0.050 | 0.773 | 0.118 | 0.663 | 0.890 | 0.907 | 0.083 | 0.880 | 0.138 | 0.677 | 0.885 |

Supplementary Table 2: Averaged precision and recall for MAGpurify, MDMcleaner, and Deepurify on $SIM_1$ simulated testing set.

The table shows that Deepurify achieves significantly higher recall values compared to MAGpurify and MDMcleaner, especially at the family, genus, and species taxonomic ranks. This implies Deepurify's effectiveness in eliminating a substantial portion of contaminated contigs at these ranks. However, it's important to note that Deepurify exhibits comparatively lower precision values at lower taxonomic ranks like family, genus, and species. This indicates that while Deepurify effectively eliminates contaminated contigs, it may unintentionally exclude a small number of core contigs.

### 5 Sequence sampling strategy during training

Inputting entire genomes into the model for training was impractical due to GPU memory limitations. Instead, we adopted a sampling strategy, wherein we randomly sampled sequences with lengths spanning 1,000 bps to 8,192 bps from genomes. We employed a Gaussian Mixture Model (GMM) consisting of ten Gaussian components to model the distribution of contig lengths. Incorporating the actual distribution of contig lengths during training was advantageous for the model to adapt to real MAGs. This GMM model was trained using the length of contigs originating from MAGs, which were binned via the MaxBin method within the $S1$ dataset. During this training phase, each sampled sequence's length exhibited a 50% probability of being the max length of 8,192 bps, with the remaining 50% probability determined through the GMM model, spanning a range from 1,000 bps to 8,192 bps.

#### 6 Sequence data augmentations

We proposed several simple data augmentation strategies for the target sequence.

- Insertions: We randomly generated a nucleotide sequence by concatenating A, T, C, and G with equal probabilities. The length of this generated sequence fell within the range of 20% to 40% of the target sequence's length, following a uniform distribution. The insertion of the generated sequence was as follows: it is either not inserted or inserted at either the beginning or end of the target sequence. These insertion options carried probabilities of 0.5, 0.25, and 0.25, respectively.

- Mutations: Each base pair in the target sequence had a 0.01 probability of being replaced by one of the other three nucleotides, with equal probabilities for each nucleotide.

- Deletions: The target sequence was either not masked or masked with 10% continuous or scattered base pairs, with probabilities of 0.5, 0.25, and 0.25, respectively.

#### 7 Sequence embedding method

We performed sequence embedding utilizing both one-hot embedding and k-mer embeddings, subsequently merging them to form a unified matrix. In the one-hot embedding procedure, we incorporated two special tokens: "$X$" denoting padding applied at the sequence's end to ensure consistent input length within a mini-batch, and "$N$" signifying representation of any unidentified characters. As a result, the one-hot embedding generated a matrix sized $L \times 6$ for each sequence, with $L$ representing the sequence's length.

We integrated k-mer embedding into our sequence embedding approach. We first created a look-up table where each row denoted a unique k-mer token and its ID. This table included two special tokens: the [PAD] token for padding sequences of varying lengths and the [UNK] token to signify any unidentified k-mer tokens within the sequences. Subsequently, we split the sequence into k-mer tokens, which are then converted to IDs through the look-up table. Lastly, a trainable matrix within the embedding layer would effectively map the ID of the k-mer token to a dense vector with $d_m(k\text{-}mer)$ dimensions. This study employed both 3-mer and 4-mer token embeddings to represent the input sequences, with dimensions set at $d_m(3\text{-}mer) = 16$ and $d_m(4\text{-}mer) = 32$, respectively.

We combined the one-hot, 3-mer, and 4-mer embeddings of the sequence with those of its reverse complement using the same embedding methods. This fusion created a unified matrix that effectively represents the input sequence. The unified matrix has a fixed dimension of $L \times 108$, with $L$ representing the sequence length. This approach provided a comprehensive and enriched representation of the input sequences, enhancing the model's robustness and effectiveness.

### 8 The architecture of modified Uniformer

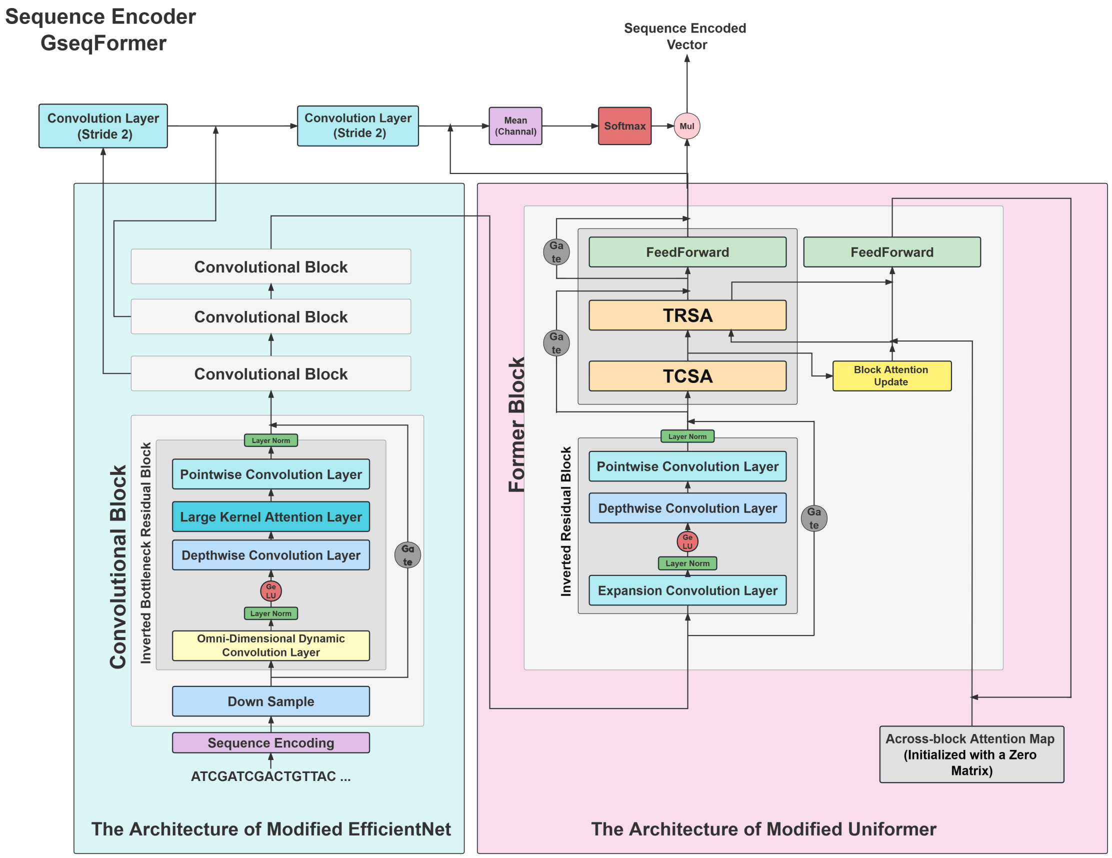


Supplementary Figure 3: The GseqFormer architecture comprises two essential components. The left segment utilizes a modified EfficientNet architecture to compress sequence length, while the right segment employs a modified Uniformer architecture with four distinct attention mechanisms designed to capture intricate taxonomic patterns within complex DNA sequences.

We substituted the Dynamic Position Embedding (DPE) blocks in Uniformer with the inverted residual blocks (IRB), which were designed from MobileNet V2 [46]. The DPE layer contained a limited number of parameters, making it less effective at handling complex sequences on its own. Therefore, we substituted it with the IRB with more trainable parameters to enhance the model's capability to process complex DNA sequences.

The Uniformer's Multi-Head Relation Aggregator (MHRA) blocks have been replaced with tensor column-wise and row-wise gated self-attention modules (TCSA, Supplementary Figure 4 and TRSA, Supplementary Figure 5), modified from the Evoformer architecture. TRSA and TCSA were designed to enhance the model's capacity to capture taxonomic patterns within complex DNA sequences by integrating four distinct attention mechanisms for their input tensor ($\psi$), which represents the sequence embedding. 1). **Embedding attention**: It evaluates the attention scores assigned to different embedding approaches for each nucleotide in the sequence. 2). **Nucleotide attention**: This mechanism captures interactions between nucleotides in a sequence, the same as the self-attention mechanism in the Transformer model. 3). **Across-block attention**: This attention mechanism links nucleotide interaction knowledge learned from different Former blocks, enabling the model to capture nucleotide interrelationships across these blocks. 4). **Spatial attention**: It extracts local spatial contexts from the attention map generated by nucleotide attention.

In both TCSA and TRSA, the input sequence embedding tensor $\psi \in \mathbf{R}^{L\times C}$ possessed two dimensions: sequence length ($L$), and channel ($C$). Initially, we partitioned the channel dimension $C$ into three equal segments in these two modules, resulting in a three-dimensional tensor denoted as $\psi_d$, with dimensions $[3, L, d_k]$, where $d_k$ is the value dividing $C$ by 3. These dimensions correspond to the number of embedding methods, the sequence length, and the channel dimension for the corresponding embedding method. This reshaped configuration allows $\psi$ to have an extra dimension to simultaneously represent three distinct embedding methods (one-hot, 3-mer, and 4-mer; refer to **Supplementary Note** 7) for a given sequence. Thus, we could apply embedding attention to this dimension.

In the TCSA module, embedding attention was applied to $\psi_d$, with a specific focus on its first dimension, representing the number of embedding approaches for a sequence. We denoted the dense vector at the $l$-th token of $\psi_d$ as $\upsilon_e \in \mathbf{R}^{3\times d_k}$. The embedding attention map $\rho_e \in \mathbf{R}^{3\times 3}$ was calculated by:

$$\rho_e = \frac{Q_e K_e^T}{\sqrt{d_k}} \quad where \quad Q_e = \upsilon_e W_e^Q, K_e = \upsilon_e W_e^K \tag{1}$$

where $W_e^Q \in \mathbf{R}^{d_k\times d_k/h}$, $W_e^K \in \mathbf{R}^{d_k\times d_k/h}$ and $h$ is the number of heads. Embedding attention effectively captured interactions among different embedding approaches within the sequence and dynamically assigned attention scores to these methods for each token in $\psi_d$.

In the TRSA module, nucleotide attention was applied to $\psi_d$, with a specific focus on its second dimension, representing the sequence length. We denoted the dense vector at the $e$-th embedding technique of $\psi_d$ as $\upsilon_n \in \mathbf{R}^{L\times d_k}$. The nucleotide attention map $\rho_n \in \mathbf{R}^{L\times L}$ was calculated by:

$$\rho_n = \frac{Q_n K_n^T}{\sqrt{d_k}}, \quad where \quad Q_n = \upsilon_n W_n^Q, K_n = \upsilon_n W_n^K$$

where $W_n^Q \in \mathbf{R}^{d_k\times d_k/h}$, $W_n^K \in \mathbf{R}^{d_k\times d_k/h}$. This attention mechanism effectively captured interactions among nucleotides within the sequence and assigned dynamic attention scores to each token in $\psi_d$.

We incorporated the across-block attention mechanism into the TRSA module to facilitate the model in establishing connections between the learned knowledge of nucleotide interactions across

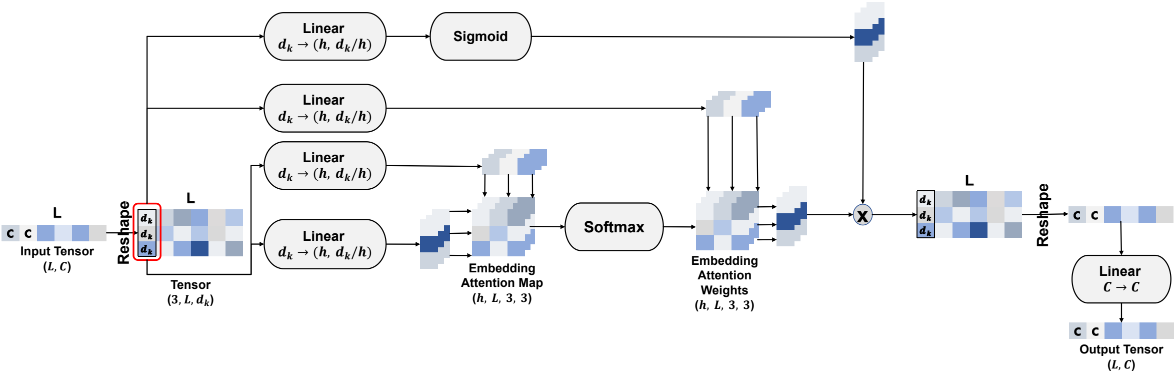


Supplementary Figure 4: The Tensor Column-wise gated Self-Attention (TCSA). Dimensions: $L$: sequence length, $C$: channels, $d_k$: channels after reshape, $h$: heads.

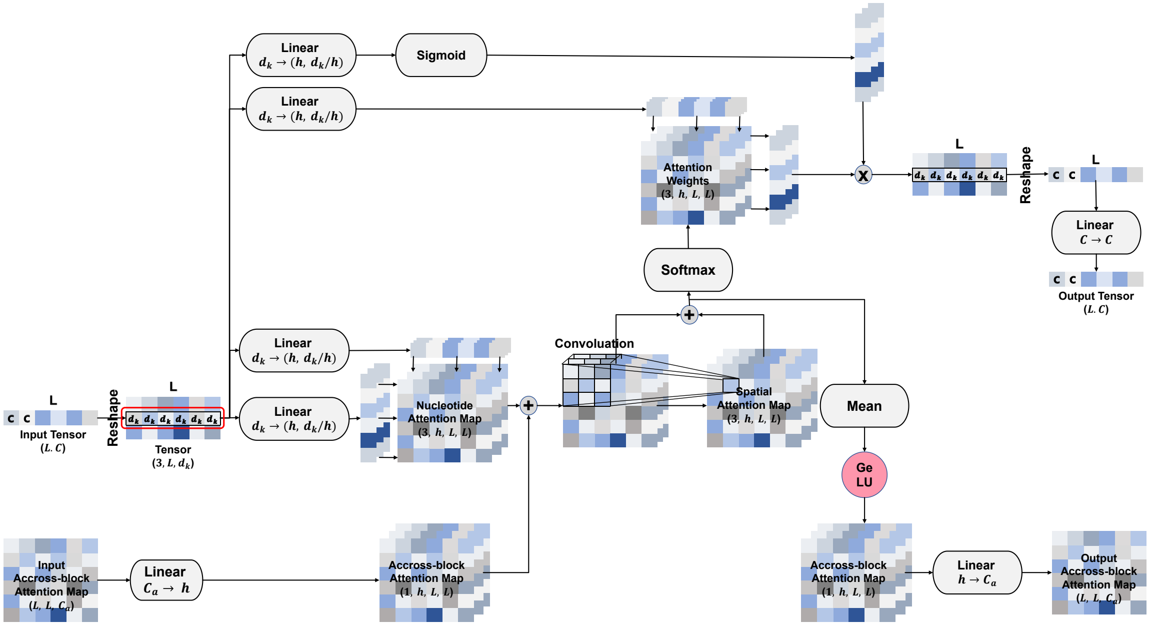


Supplementary Figure 5: The Tensor Row-wise gated Self-Attention (TRSA). Dimensions: $L$: sequence length, $C$: channels for tokens, $d_k$: channels for tokens after reshape, $C_a$: channels for across-block attention map, $h$: heads.

different blocks. Initially, the across-attention map ($\rho_a \in \mathbf{R}^{L \times L \times C_a}$) was set as a zero matrix. $\rho_a$ was then updated (added) through the block attention update (BAU) module (**Supplementary Figure** 6) based on the TCSA's output. Subsequently, the updated $\rho_a$ was fed into TRSA. After a linear transformation, it was added with $\rho_n$ to serve as an additional nucleotide interaction attention map derived from previous blocks. Further refinement of $\rho_a$ was achieved through a subsequent feedforward module. The implementation of the across-block attention mechanism facilitates the sharing and propagation of nucleotide interaction knowledge learned within individual blocks across different parts of the model.

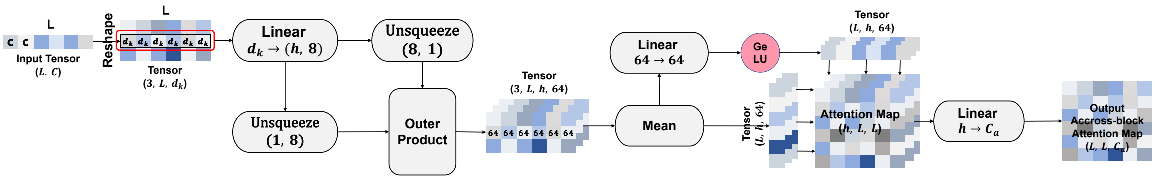


Supplementary Figure 6: The Block Attention Update (BAU) module. Dimensions: $L$: sequence length, $C$: channels for tokens, $d_k$: channels for tokens after reshape, $C_a$: channels for across-block attention map, $h$: heads.

Transformer-based models typically excel at capturing the global context of a sequence but may be limited in capturing its local context [47]. Hence, we extended the nucleotide attention map and across-block attention map by incorporating spatial attention into it. The spatial attention was employed to extract local spatial contexts from $\rho_n + Linear(\rho_a)$ by treating it as an image through the application of a convolutional layer. This process resulted in the generation of a spatial attention map denoted as $\rho_s$. In the TRSA module, $\rho_s$ was incorporated into $\rho_n + Linear(\rho_a)$ as a local context attention map. The spatial attention improved the model's capacity to capture and comprehend the local context within a given sequence.

#### 9 The architecture of modified EfficientNet

We modified EfficientNet [38] by incorporating three changes. 1). The expansion convolutional layers present in inverted bottleneck residual blocks (IBRB) were substituted with omni-dimensional dynamic convolution [39], which has been demonstrated to yield superior accuracy in comparison to static convolutions [39]. 2). Substituted the squeeze and excitation blocks [48] in IBRB with the large kernel attention layers [40] to calculate the token-wise and channel-wise attention scores for sequence embeddings. 3). DeepNorm [41] was incorporated to prevent gradient vanishing during model training.

#### 10 Strategy for generating negative taxonomic lineages during training

We denoted $T_{k_i} = [\leqslant t_{k_i}]$ as the prefix of taxonomic lineage $T_i$ before $k_i$ rank, where $k_i \in \{phylum, class, order, family, genus, species\}$. During training, $k_i$ would be randomly selected and we treat $T_{k_i}$ as the positive taxonomic lineage for sequence $s_i$.

We denoted the negative taxonomic lineage as $T_{k_j}$. During contrastive training, we would select 99 negative taxonomic lineages from the taxonomic tree. The half-number of negative taxonomic lineages have three characteristics: 1. $k_i = k_j$, 2. $[< t_{k_i}] = [< t_{k_j}]$, 3. $t_{k_i} \neq t_{k_j}$. The remaining half of these negative lineages would be randomly drawn from the taxonomic tree, ensuring they are distinct with $T_{k_i}$. This generation method is pivotal in Deepurify's contrastive training process, where $T_{k_j}$ with the three specified characteristics act as hard negative samples, increasing the contrastive training difficulty and enhancing the model's performance [49].

#### 11 Solve the imbalance of phyla labels issue in $GS_c$ and $GS_p$

In our data processing pipeline, we incorporated an oversampling strategy to mitigate the issue of imbalanced phyla labels observed in $GS_c$ and $GS_p$. This imbalance problem was caused by the varying number of species present across different phyla. The oversampling approach involved duplicating genomes within a phylum until the total number of genomes from that phylum reached a predefined threshold, specifically a minimum of 500 for $GS_c$ and 20 for $GS_p$. This strategy was only applied to the phylum containing fewer than 500 or 20 species, respectively.

Furthermore, we adopted the focal loss [42] in the training stage to address the issue of label imbalance in the other taxonomic ranks. This additional measure enhanced our model's robustness to imbalanced data distributions.

#### 12 Hyper-parameter setting for training

We applied the hyper-parameter for training Deepurify, similar to the training configuration of Uniformer. Specifically, we established the stochastic depth rate [50] at 0.1 for the EfficientNet and set the dropout rate [51] to 0.15. We set the weight decay, learning rate, batch size, and number of negative lineages for contrastive learning as $5e^{-4}$, $1e^{-4}$, 16, and 99, respectively. Training Deepurify involved utilizing the AdamW optimizer [52] in conjunction with a cosine learning rate schedule [53], spanning a training period of 128 epochs. The initial 10 epochs were used for linear warm-up.

We fine-tuned Deepurify subsequently to minimize the influence of homologous sequences during training. The basic parameter would not change besides setting 0 for stochastic depth rate and dropout. The taxonomic encoder (LSTM) parameter was fixed, and there were only 15 epochs for fine-tuning.

Only $L_{ST}$ and $L_{PR}$ would be calculated during fine-tuning, and $L_{ST}$ was replaced with cross-entropy loss but not focal loss. To mitigate the impact of homologous sequences on the model, we computed the loss $L_{ST}$ for a given sequence $s_i$ only within the context of contrastive training if the absolute difference between the top-1 and top-2 predicted probabilities of lineages exceeded 0.05.

#### 13 Removing duplicated SCGs within a node

We identified SCGs using Prodigal and the HMM tool, resulting in a comprehensive list that contained contigs to their corresponding SCGs.We then arranged these contigs in descending order of length and assigned them to sets one by one, while simultaneously recording the SCGs in sets. We created a new set for a contig if we encountered duplicate SCGs within any set while adding that contig, This iterative process continued until all contigs found their place in different sets, resulting in multiple divisions.

#### 14 Determining the threshold of total contig length to remove low-quality MAGs

We used all MAGs from the IBS-D cohort and assessed their quality with CheckM. We then calculated the total contig lengths of all MAGs and plotted their distribution for low-quality, medium-quality, and high-quality MAGs, respectively (**Supplementary Figure** 7).

Our analysis showed significant differences in the distributions for MAGs with different qualities. If total lengths of MAGs were shorter than 550kbps, We found 43% MAGs were low-quality, $< 1\%$ MAGs were medium-quality and no MAG was classified high-quality.

#### 15 Traverse the MAG-separated tree

Deepurify applied post-order traversal to traverse the MAG-separated tree to maximize the total number of medium- and high-quality MAGs. It maintained a list for each node to store its selected child nodes. Initially, the leaf nodes' lists only contained the nodes themselves, while the lists of the other nodes were empty. We referred to the node currently being traversed as $V_C$. $V_{M\_}$ and $V_{H\_}$ represented the Medium-quality and High-quality divisions in the lists of all child nodes of $V_C$. The node comparison and selection algorithm was presented in Algorithm 1. The comparison and selection process occurs recursively, starting from the left nodes and progressing up to the root node. Ultimately, this process results in a collection of nodes stored in the list of the root node.

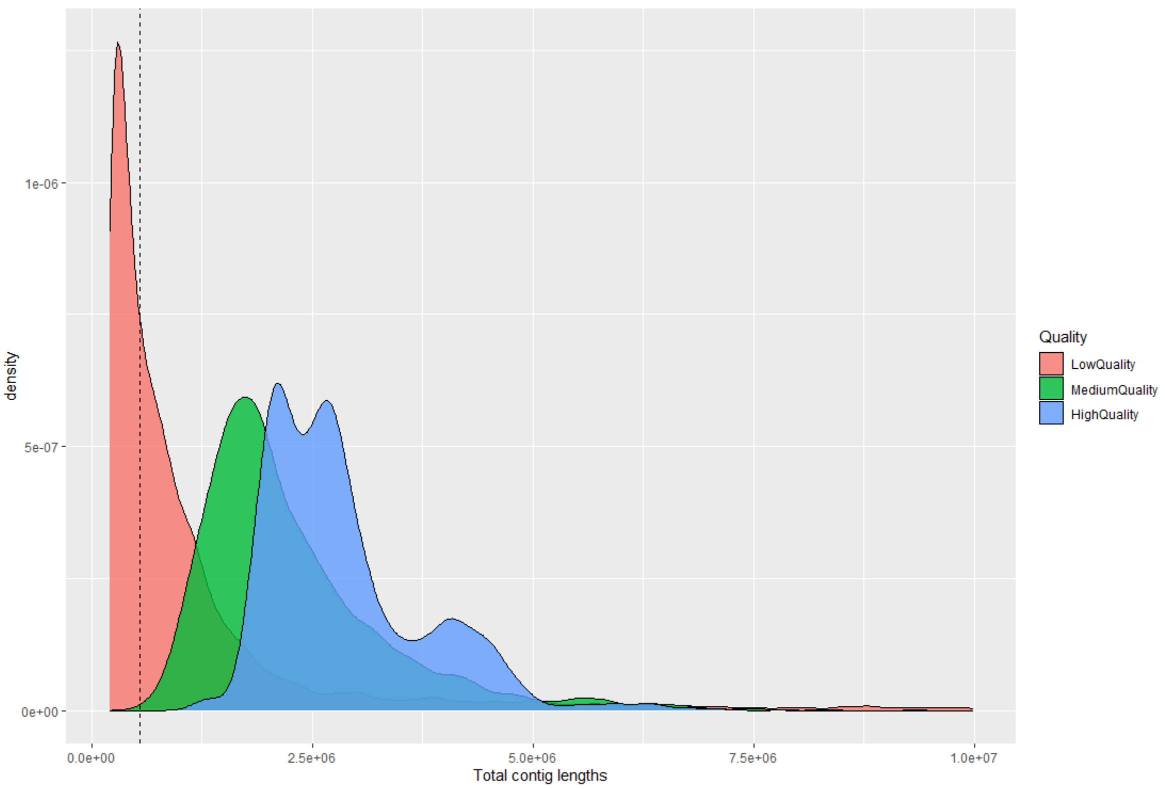


Supplementary Figure 7: The distribution of total contig lengths among low-, medium-, and high-quality MAGs.

**Algorithm 1** The comparison and selection of nodes

**Input**: A list contains medium-quality nodes: $L_M = [V_{M_1}, ..., V_{M_m}]$,
**Input**: A list contains high-quality nodes: $L_H = [V_{H_1}, ..., V_{H_h}]$,
**Input**: The node that DFS is currently traversing: $V_C$.
**Output**: A list contains the nodes that need to be preserved.
**function** comparison($L_M$, $L_H$, $V_C$):
    res = [$L_M$, $L_H$]
    $h$ = size($L_H$)
    $m$ = size($L_M$)
    if $h$ == 0:
        if $V_C$.quality == "HighQuality":
            res = [$V_C$]
        elif $V_C$.quality == "MediumQuality":
            if $m$ == 0:
                res = [$V_C$]
            elif $m$ == 1:
                if QS($V_C$) ¿ QS(res[0]): ▷ QS is a function to calculate quality score of node
                    res = [$V_C$]
    elif $h$==1 and $m$ == 0 and $V_C$.quality == "HighQuality":
        if QS($V_C$) ¿ QS(res[0]):
            res = [$V_C$]
    return res

#### 16 Software versions and computational environment

Our study employed MAGpurify (v2.1.2) and MDMcleaner (v0.8.3) for MAG decontamination. The development of Deepurify involved the utilization of Python (v3.11.1) along with PyTorch (v2.0.1 + cu118). The SCGs calling was executed using Prodigal (v2.6.3) and HMMER (v3.3.1). The computation of the balanced macro F1-score was performed using Scikit-Learn (v1.2.0). The evaluation of MAG quality was carried out using CheckM (v1.2.2). For binning, we employed MaxBin (v2.2.7), VAMB (v3.0.3), MetaBAT2 (v2.12.1), and CONCOCT (v1.1.0). The annotation of MAGs was executed using GTDB-Tk (v1.4.0). In this study, metaSPAdes (v3.15.0) was employed for assembly. For sequence alignment, BLASTN (v2.9.0+) was utilized.

The experiments were conducted on a compute node with an AMD EPYC 7542 processor comprising 32 cores (64 threads) and 256 GB of memory. To expedite the Deepurify training, four NVIDIA Tesla V100 32GB GPUs were engaged. During inference, the utilization was limited to two GTX 3090 24GB GPUs, with two threads allocated for data feeding to each GPU. Other MAG decontamination tools were run with four threads.

### Supplementary Figures

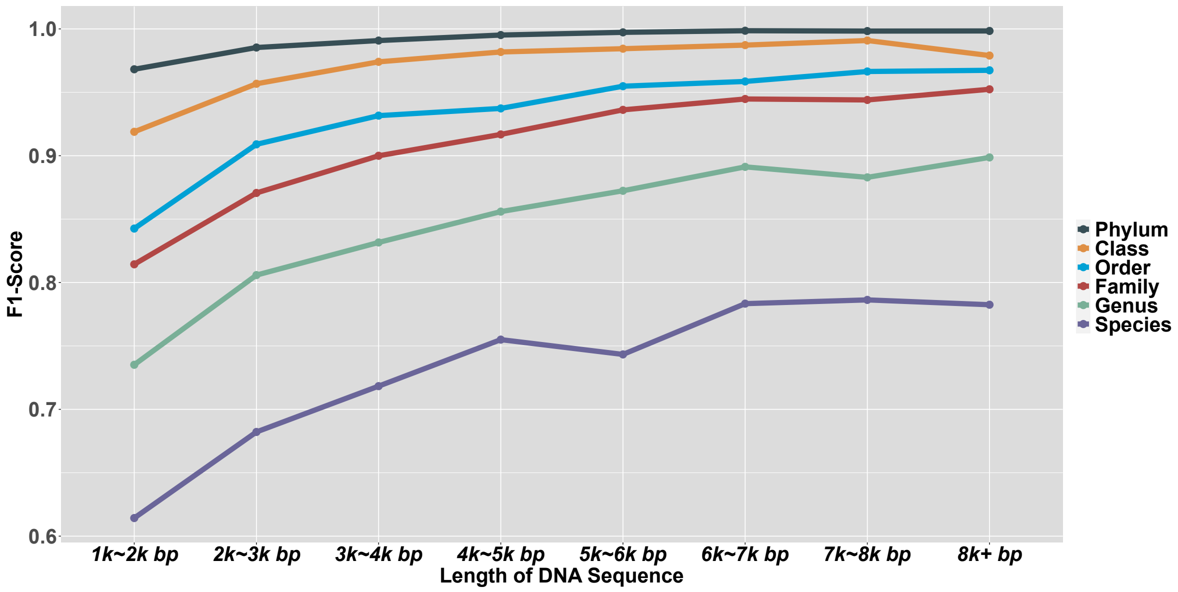


Supplementary Figure 8: The averaged macro F1-score for different length categories with different taxonomic ranks for Deepurify.

### Supplementary Tables

| Taxonomy | Contamination 5% | | | Contamination 10% | | |
|---|---|---|---|---|---|---|
| | MAGpurify | MDMcleaner | Deepurify | MAGpurify | MDMcleaner | Deepurify |
| Phylum | 0.742 | 0.988 | 0.988 | 0.738 | 0.995 | 0.993 |
| Class | 0.555 | 0.994 | 0.969 | 0.589 | 0.990 | 0.969 |
| Order | 0.523 | 0.960 | 0.936 | 0.539 | 0.976 | 0.932 |
| Family | 0.460 | 0.802 | 0.889 | 0.509 | 0.811 | 0.908 |
| Genus | 0.416 | 0.655 | 0.859 | 0.463 | 0.529 | 0.843 |
| Species | 0.429 | 0.476 | 0.758 | 0.4071 | 0.450 | 0.741 |
| | Contamination 15% | | | Contamination 20% | | |
| Taxonomy | MAGpurify | MDMcleaner | Deepurify | MAGpurify | MDMcleaner | Deepurify |
| Phylum | 0.641 | 0.997 | 0.992 | 0.608 | 0.996 | 0.989 |
| Class | 0.505 | 0.981 | 0.974 | 0.548 | 0.987 | 0.972 |
| Order | 0.503 | 0.972 | 0.916 | 0.494 | 0.975 | 0.936 |
| Family | 0.454 | 0.857 | 0.901 | 0.481 | 0.865 | 0.920 |
| Genus | 0.458 | 0.554 | 0.846 | 0.430 | 0.657 | 0.821 |
| Species | 0.383 | 0.413 | 0.691 | 0.404 | 0.426 | 0.702 |

Supplementary Table 3: Averaged balanced macro F1-scores for MAGpurify, MDMcleaner and Deepurify in $SIM_1$.

| Taxonomy | Contamination 5% | | | | Contamination 10% | | | |
|---|---|---|---|---|---|---|---|---|
| | $SIM_2^1$ | $SIM_2^2$ | $SIM_2^3$ | $SIM_2^4$ | $SIM_2^1$ | $SIM_2^2$ | $SIM_2^3$ | $SIM_2^4$ |
| Phylum | 0.994 | 0.986 | 0.785 | 0.783 | 0.992 | 0.980 | 0.836 | 0.799 |
| Class | 0.986 | 0.971 | 0.818 | 0.789 | 0.986 | 0.978 | 0.790 | 0.797 |
| Order | 0.957 | 0.925 | 0.807 | 0.774 | 0.978 | 0.943 | 0.798 | 0.742 |
| Family | 0.940 | 0.872 | 0.750 | 0.743 | 0.935 | 0.897 | 0.738 | 0.744 |
| Genus | 0.946 | 0.798 | 0.741 | 0.682 | 0.937 | 0.799 | 0.707 | 0.664 |
| Species | 0.778 | 0.687 | 0.633 | 0.608 | 0.792 | 0.607 | 0.604 | 0.599 |
| | Contamination 15% | | | | Contamination 20% | | | |
| Taxonomy | $SIM_2^1$ | $SIM_2^2$ | $SIM_2^3$ | $SIM_2^4$ | $SIM_2^1$ | $SIM_2^2$ | $SIM_2^3$ | $SIM_2^4$ |
| Phylum | 0.994 | 0.980 | 0.860 | 0.778 | 0.992 | 0.983 | 0.837 | 0.818 |
| Class | 0.984 | 0.951 | 0.816 | 0.802 | 0.985 | 0.971 | 0.728 | 0.783 |
| Order | 0.973 | 0.926 | 0.763 | 0.758 | 0.973 | 0.927 | 0.771 | 0.727 |
| Family | 0.956 | 0.883 | 0.717 | 0.759 | 0.949 | 0.869 | 0.744 | 0.724 |
| Genus | 0.935 | 0.835 | 0.730 | 0.674 | 0.940 | 0.882 | 0.702 | 0.700 |
| Species | 0.795 | 0.664 | 0.563 | 0.592 | 0.795 | 0.661 | 0.620 | 0.589 |

Supplementary Table 4: Averaged balanced macro F1-score of Deepurify for four permuted scenarios for $SIM_2$.

| Taxonomic Lineages | | | | | |
|---|---|---|---|---|---|
| Phylum | Class | Order | Family | Genus | Species |
| Firmicutes | Bacilli | RF39 | UBA660 | CAG-914 | CAG-914 sp000437895 |
| Firmicutes | Bacilli | RF39 | UBA660 | UBA5026 | UBA5026 sp900552335 |
| Actinobacteria | Coriobacteriia | Coriobacteriales | Coriobacteriaceae | Collinsella | Collinsella sp900541055 |
| Firmicutes | Bacilli | Staphylococcales | Gemellaceae | Gemella | Gemella morbillorum |
| Firmicutes | Bacilli | RF39 | UBA660 | CAG-460 | CAG-460 sp900551525 |

Supplementary Table 5: Five new species identified after purification for IBS cohort. Collinsella sp900541055 has a significant P-value after association analysis

| Taxonomic Lineages | | | | |
|---|---|---|---|---|
| Phylum | Class | Order | Family | Genus |
| Synergistota | Synergistia | Synergistales | Synergistaceae | Cloacibacillus |

Supplementary Table 6: One new genus identified after purification for IBS cohort.

| Taxonomy \ Length (bps) | 1000 - 2000 | 2000 - 3000 | 3000 - 4000 | 4000 - 5000 | 5000 - 6000 | 6000 - 7000 | 7000 - 8000 | 8000 - 8192 |
|---|---|---|---|---|---|---|---|---|
| Phylum | 0.968 | 0.985 | 0.990 | 0.995 | 0.997 | 0.998 | 0.998 | 0.998 |
| Class | 0.918 | 0.956 | 0.974 | 0.981 | 0.984 | 0.987 | 0.991 | 0.978 |
| Order | 0.842 | 0.908 | 0.931 | 0.937 | 0.954 | 0.958 | 0.966 | 0.967 |
| Family | 0.814 | 0.871 | 0.899 | 0.916 | 0.936 | 0.944 | 0.943 | 0.952 |
| Genus | 0.735 | 0.805 | 0.831 | 0.855 | 0.872 | 0.891 | 0.882 | 0.898 |
| Species | 0.614 | 0.682 | 0.718 | 0.754 | 0.743 | 0.783 | 0.786 | 0.782 |

Supplementary Table 7: Averaged balanced macro F1-score for length categories with different taxonomic ranks for Deepurify.

| ***CONCOCT*** | No Purification | | MAGpurify | | MDMcleaner | | Deepurify | |
|---|---|---|---|---|---|---|---|---|
| Dataset | High | Medium | High | Medium | High | Medium | High | Medium |
| CAMI I Medium 1 | 28 | 9 | 30 | 7 | 29 | 9 | 30 | 11 |
| CAMI I Medium 2 | 34 | 15 | 33 | 15 | 34 | 15 | 36 | 19 |
| CAMI I Low | 11 | 5 | 12 | 4 | 11 | 4 | 11 | 7 |
| CAMI I High | 29 | 29 | 26 | 32 | 43 | 44 | 59 | 120 |
| $S_1$ | 36 | 18 | 29 | 26 | 36 | 21 | 37 | 53 |

| ***MetaBAT2*** | No Purification | | MAGpurify | | MDMcleaner | | Deepurify | |
|---|---|---|---|---|---|---|---|---|
| Dataset | High | Medium | High | Medium | High | Medium | High | Medium |
| CAMI I Medium 1 | 25 | 12 | 23 | 10 | 27 | 10 | 26 | 13 |
| CAMI I Medium 2 | 39 | 13 | 38 | 14 | 37 | 16 | 39 | 14 |
| CAMI I Low | 13 | 6 | 13 | 6 | 12 | 7 | 13 | 7 |
| CAMI I High | 115 | 112 | 105 | 118 | 116 | 117 | 116 | 139 |
| $S_1$ | 50 | 58 | 37 | 68 | 50 | 58 | 50 | 65 |

| ***MaxBin*** | No Purification | | MAGpurify | | MDMcleaner | | Deepurify | |
|---|---|---|---|---|---|---|---|---|
| Dataset | High | Medium | High | Medium | High | Medium | High | Medium |
| CAMI I Medium 1 | 19 | 5 | 22 | 4 | 25 | 10 | 25 | 14 |
| CAMI I Medium 2 | 28 | 15 | 29 | 15 | 34 | 18 | 33 | 20 |
| CAMI I Low | 11 | 4 | 11 | 6 | 13 | 6 | 12 | 7 |
| CAMI I High | 62 | 61 | 67 | 74 | 80 | 85 | 75 | 119 |
| $S_1$ | 31 | 28 | 29 | 37 | 44 | 34 | 38 | 63 |

| ***VAMB*** | No Purification | | MAGpurify | | MDMcleaner | | Deepurify | |
|---|---|---|---|---|---|---|---|---|
| Dataset | High | Medium | High | Medium | High | Medium | High | Medium |
| CAMI I Medium 1 | 25 | 9 | 25 | 9 | 25 | 9 | 25 | 9 |
| CAMI I Medium 2 | 27 | 7 | 27 | 8 | 27 | 9 | 29 | 11 |
| CAMI I Low | 11 | 4 | 11 | 4 | 10 | 5 | 11 | 5 |
| CAMI I High | 99 | 56 | 94 | 62 | 98 | 58 | 98 | 67 |
| $S_1$ | 30 | 25 | 24 | 31 | 30 | 25 | 32 | 30 |

Supplementary Table 8: The number of high- and medium-quality MAGs in five public datasets before purification (No Purification), and after MAGpurify, MDMcleaner, and Deepurify purification.

| ***CONCOCT*** | No Purification | MAGpurify | MDMcleaner | Deepurify |
|---|---|---|---|---|
| CAMI I Medium 1 | 3041.33 | 3102.04 | 3234.65 | 3339.61 |
| CAMI I Medium 2 | 3835.34 | 3857.59 | 4035.47 | 4331.21 |
| CAMI I Low | 1315.76 | 1349.11 | 1297.6 | 1445.22 |
| CAMI I High | 4385.38 | 4422.13 | 6596.01 | 11329.95 |
| $S_1$ | 4473.11 | 4466.04 | 4709.28 | 5668.46 |
| ***MetaBAT2*** | No Purification | MAGpurify | MDMcleaner | Deepurify |
| CAMI I Medium 1 | 3076.61 | 2762.12 | 3127.94 | 3175.27 |
| CAMI I Medium 2 | 4316.47 | 3893.56 | 4426.45 | 4412.11 |
| CAMI I Low | 1635.05 | 1630.92 | 1604.62 | 1695.48 |
| CAMI I High | 16949.9 | 16855.01 | 17521.99 | 18200.68 |
| $S_1$ | 7999.76 | 7930.49 | 8079.96 | 8268.43 |
| ***MaxBin*** | No Purification | MAGpurify | MDMcleaner | Deepurify |
| CAMI I Medium 1 | 2083.78 | 2315.52 | 2959.23 | 3075.27 |
| CAMI I Medium 2 | 3474.14 | 3662.42 | 4187.73 | 4159.03 |
| CAMI I Low | 1253.52 | 1434.82 | 1620.37 | 1601.22 |
| CAMI I High | 8908.38 | 10208.37 | 12568.25 | 13449.79 |
| $S_1$ | 4369.67 | 4864.69 | 5885.65 | 6519.08 |
| ***VAMB*** | No Purification | MAGpurify | MDMcleaner | Deepurify |
| CAMI I Medium 1 | 2863.67 | 2872.31 | 2893.24 | 2879.4 |
| CAMI I Medium 2 | 2973.75 | 3059.16 | 3138.38 | 3393.77 |
| CAMI I Low | 1268.58 | 1271.57 | 1258.94 | 1307.29 |
| CAMI I High | 12871.03 | 12886.54 | 12934.18 | 13327.5 |
| $S_1$ | 4586.81 | 4563.97 | 4595.11 | 4960.31 |

Supplementary Table 9: The quality scores in five public datasets before purification (No Purification), and after MAGpurify, MDMcleaner, and Deepurify purification.

| ***The 37 Phyla in the Training Set*** $GS_c$ | | | | |
|---|---|---|---|---|
| Acidobacteria | Actinobacteria | Aquificae | Bacteroidetes | Candidatus-Gottesmanbacteria |
| Candidatus-Kaiserbacteria | Candidatus-Levybacteria | Candidatus-Magasanikbacteria | Candidatus-Moranbacteria | Candidatus-Nomurabacteria |
| Candidatus-Omnitrophica | Candidatus-Parcubacteria | Candidatus-Peregrinibacteria | Candidatus-Roizmanbacteria | Candidatus-Saccharibacteria |
| Candidatus-Uhrbacteria | Candidatus-Woesebacteria | Candidatus-Yanofskybacteria | Chlamydiae | Chlorobi |
| Chloroflexi | Crenarchaeota | Cyanobacteria | Deinococcus-Thermus | Elusimicrobia |
| Euryarchaeota | Firmicutes | Fusobacteria | Nitrospirae | Planctomycetes |
| Proteobacteria | Spirochaetes | Synergistetes | Tenericutes | Thaumarchaeota |
| Thermotogae | Verrucomicrobia | | | |

Supplementary Table 10: $GS_c$ training set includes the genomes from 37 phyla.

| ***The 12 Phyla in the Training Set*** $GS_p$ | | | | |
|---|---|---|---|---|
| Bacteroidetes | Candidatus-Saccharibacteria | Crenarchaeota | Cyanobacteria | Deferribacteres |
| Deinococcus-Thermus | Euryarchaeota | Firmicutes | Fusobacteria | Planctomycetes |
| Proteobacteria | Balneolaeota | | | |

Supplementary Table 11: $GS_p$ training set includes the genomes from 12 phyla.
